# Sleep pressure is differentially regulated by molecularly distinct subtypes of Lhx6-positive and Lhx6-negative neurons of the zona incerta

**DOI:** 10.1101/2025.07.30.667705

**Authors:** Parris Washington Chandler, Sang Soo Lee, Leighton H. Duncan, Dong Won Kim, Mark N. Wu, Seth Blackshaw

## Abstract

Sleep pressure, the accumulating drive towards sleep during wakefulness, is shaped by Lhx6-positive GABAergic neurons in the zona incerta (ZI). Here, we show that these neurons are broadly activated both by spontaneous and experimentally-elevated sleep pressure, and remain active for hours into recovery sleep. Activation is stronger in anterior ZI and varies across molecularly defined subgroups: Nkx2-2-positive neurons respond vigorously, whereas Calb2-positive neurons respond weakly. We also identify Lhx6-negative/Slc32a1-positive GABAergic ZI neurons with distinct sleep pressure responses. Selective genetic ablation of *Nkx2-2* in Lhx6-positive neurons reduces the number of Lhx6-positive neurons, alters their distribution, blunts sleep pressure-induced activation of both Lhx6-positive and negative cells, and increases sleep duration. These findings show that developmentally specified, molecularly heterogeneous Lhx6-positive ZI neurons form a key hub for regulating sleep homeostasis, and offer new insight into the circuitry that controls sleep pressure.

## Introduction

Sleep is an evolutionarily-conserved state that is essential for the survival of all organisms examined to date (Keene and Duboue 2018; Freiberg 2020; Miyazaki, Liu, and Hayashi 2017). In recent years, multiple neuronal subtypes that rapidly regulate bistable sleep-wake transitions have been identified (Eban-Rothschild, Appelbaum, and de Lecea 2018; Xu et al. 2015; Yu et al. 2019; Vanini and Torterolo 2021). The drive to sleep is controlled by an activity/time-dependent component referred to as sleep pressure, that progressively accumulates during wakefulness and dissipates during sleep, as well as by circadian factors (Skeldon and Dijk 2025; Borbély 2022). However, the molecular basis of this activity/time-dependent drive, and thus of sleep homeostasis, remains unknown (Thomas et al. 2020; Duhart, Inami, and Koh 2023).

Recent evidence from both Drosophila and mammals has suggested that activity-dependent changes in rapid-acting wake and sleep-promoting neurons may drive the gradual accumulation of sleep pressure (Sawada et al. 2024; Thomas et al. 2020). However, studies in mice have identified discrete neuronal subpopulations that are activated selectively in response to both natural or induced changes in sleep pressure, and are distinct from those controlling rapid sleep-wake transitions. One such neuronal population is Lhx6-positive GABAergic neurons of the zona incerta (ZI) (Liu et al. 2017a; Lee et al. 2025; Kim et al. 2021a), a small yet highly complex region that broadly regulates sensory integration, innate behaviors, and both motivational and emotional states (Mitrofanis 2005; Venkataraman et al. 2019; Li, Rizzi, and Tan 2021; Wang et al. 2020; Monosov et al. 2022; X. Zhang and van den Pol 2017). Although these neurons are immediately presynaptic to several arousal-promoting cell types, their chemogenetic activation induces sleep with notably delayed and prolonged kinetics, ranging from 2-8 hours following CNO administration (Liu et al. 2017a). Recent work has shown that glutamatergic neurons of the thalamic reuniens nucleus (ReN), which are both selectively responsive to induced sleep deprivation and project to anterior Lhx6-positive ZI neurons, promote sleep with similar delayed kinetics (Lee et al. 2025). Elevated sleep pressure both structurally and functionally strengthens the synaptic connection between ReN and Lhx6-positive ZI neurons, and Lhx6-positive neurons are essential for the sleep-promoting function of ReN neurons (Lee et al. 2025). Together, these findings suggest that this ReN-ZI connection forms a dedicated neural circuit for sensing and conveying sleep pressure, distinct from the circuitry that mediate rapid bistable sleep-wake transitions (Han et al. 2014; Kroeger et al. 2018; Kashiwagi et al. 2024).

This raises the question of exactly how sleep pressure is regulated by Lhx6-positive ZI neurons. Previous studies indicate substantial functional heterogeneity within this population. Using Fos expression as a readout for activity, at least 25% of Lhx6-positive ZI neurons are active even in the early evening, when sleep pressure is the lowest, while many remain inactive even under high levels of experimentally-induced sleep pressure (Liu et al. 2017a). These neurons also remain active even during early stages of recovery sleep (Liu et al. 2017a). However, it is not known whether the same subpopulations respond to natural versus induced sleep pressure, or during recovery sleep.. Other studies have demonstrated substantial molecular heterogeneity among these cells (Kim et al. 2021a, 2020, 2025). This raises the possibility that molecularly distinct Lhx6-positive ZI subtypes may serve different functions in sleep homeostasis.

In this study, we systematically applied multiplexed single-molecule fISH (smfISH) analysis for *Lhx6*, *Fos*, and molecular markers that define distinct subtypes of Lhx6-positive ZI neurons. We confirm that these neurons are broadly activated by both naturally occurring and sleep deprivation-induced increases in sleep pressure, and remain strongly activated for at least three hours after the onset of recovery sleep. Anterior *Lhx6* mRNA-positive neurons show relatively stronger sleep pressure-induced activation, broadly coinciding with patterns of ReN innervation. Distinct patterns of *Fos* mRNA induction were observed in molecularly distinct subpopulations of *Lhx6*-positive ZI neurons, with *Nkx2-2*, *Nfia*, and *Calb1* mRNA-positive cells showing the strongest responses to sleep pressure, whereas *Calb2*-positive cells respond weakly. We also identified subpopulations of *Lhx6*-negative, *Slc32a1*-positive GABAergic ZI neurons that were differentially activated by sleep pressure. Finally, using intersectional genetics, we show that loss of function of the homeodomain transcription factor *Nkx2-2* markedly reduces the number of Lhx6-positive neurons, redistributes them along the anterior-posterior axis, diminishes their sleep-pressure-induced activity, and increases both daytime and nighttime sleep. These results indicate that *Lhx6*-positive ZI neurons play a central but complex role in regulating sleep homeostasis.

## Results

### Multiplexed smfISH analysis of *Lhx6*-positive ZI neurons

To map spatial, temporal, and subtype-dependent activation of *Lhx6*-positive ZI neurons, we used the HiPlex platform (ACD Bio) to simultaneously profile up to 12 different genes, under a range of both naturally occurring and induced elevated sleep pressure. Core probes targeted *Lhx6* and *Fos*, as well as *Slc32a1*, which broadly labels GABAergic neurons in the ZI (Watson, Lind, and Thomas 2014; Li, Rizzi, and Tan 2021). Additional probes marked subtypes previously defined in the ZI (Kim et al. 2021a; Watson, Lind, and Thomas 2014; Zhao et al. 2019; M. Zhang et al. 2023; Mitrofanis 2005): *Nkx2-2*, *Calb1*, *Calb2*, *Nfia*, *Cck*, *Penk*, *Pvalb*, *Gal*, *Nos1*, and *Pnoc*. *Slc17a6* was used in some studies to detect glutamatergic neurons. Pilot analyses showed that *Nkx2-2*, *Calb1*, *Calb2*, *Nfia*, and *Cck* gave the strongest and most robust signal, and these were used for subsequent analysis (Table 1).

**Table 1:** Hiplex probes used in the study. Probes used in each iteration of the Hiplex analysis. *Nfix, Nos1*, and *Pnoc* were removed due to low, inconsistent signals. *Pvalb, Penk*, and *Gal* were not included in the final analysis because of data loss due to tissue damage (TD) by the 3^rd^ round of staining. *Slc17a6* was only in a small subset of experiments and was not included in the final analysis.

| Round/Channel | Fluorophore | 1<sup>st</sup> Iteration | 2<sup>nd</sup> Iteration | 3<sup>rd</sup> Iteration |
| --- | --- | --- | --- | --- |
| R1:T1 | 488 (Green) | Lhx6<br>In Data | Lhx6<br>In Data | Lhx6<br>In Data |
| R1:T2 | 550 (Orange) | Nkx2-2<br>In Data | Fos<br>In Data | Nkx2-2<br>In Data |
| R1:T3 | 647N (Far Red) | Nfia<br>In Data | Nkx2-2<br>In Data | Nfia<br>In Data |
| R1:T4 | 750 (Near Red) | Calb1<br>In Data | Slc17a6<br>Not in data analysis | Calb1<br>In Data |
| R1:T5 | 488 (Green) | Calb2<br>In Data | Slc32a1<br>In Data | Calb2<br>In Data |
| R2:T6 | 550 (Orange) | Nfix<br>Removed | Nfia<br>In Data | Slc32a1<br>In Data |
| R2:T7 | 647N (Far Red) | Fos<br>In Data | Nfix<br>Removed | Fos<br>In Data |
| R2:T8 | 750 (Near Red) | Cck<br>In Data | Cck<br>In Data | Cck<br>In Data |
| R3:T9 | 488 (Green) | Nos1<br>Removed | Nos1<br>Removed | Nos1<br>Removed |
| R3:T10 | 550 (Orange) | Pnoc<br>Removed | Pvalb<br>TD* | Pvalb<br>TD |
| R3:T11 | 647N (Far Red) | Penk<br>TD | Penk<br>TD | Penk<br>TD |
| R3:T12 | 750 (Near Red) | Gal<br>TD | Gal<br>TD | Gal<br>TD |

\*TD - Samples incurred severe tissue damage by R3 of imaging rendering foci uncountable

Using these probe sets, we first analyzed the distribution of *Lhx6*-positive cells along the anterior-posterior axis of the ZI at Zeitgeber Time 6 (ZT6), which corresponds to 6 hours following lights on, which is a mild to moderate sleep pressure state in nocturnal mice (Fig. 1A, Table 2) (Mitler et al. 1977). *Lhx6* mRNA-positive cells comprised 35.2% of all ZI cells (Fig. 1B), consistent with estimates from Lhx6 immunohistochemistry and Lhx6-eGFP reporter lines (Liu et al. 2017a; Kim et al. 2021a). Most of these cells (33.7% of all ZI cells) showed relatively low levels of *Lhx6* expression, ranging from 1-5 puncta associated with each DAPI-positive cell (Fig. 1A, B, Table 2) , whereas only 1.5 % showed high expression. The density of *Lhx6*-positive cells decreased from anterior to posterior ZI (Fig. 1C): anterior (-0.955 to -1.554 mm Bregma) 2.7 % high / 53.2 % low, medial (-1.555 to -2.154 mm Bregma) 1.5 % / 31 %, and posterior (-2.155 to - 2.780 mm Bregma) 0.8 % / 21.8 %.

**Figure 1:**
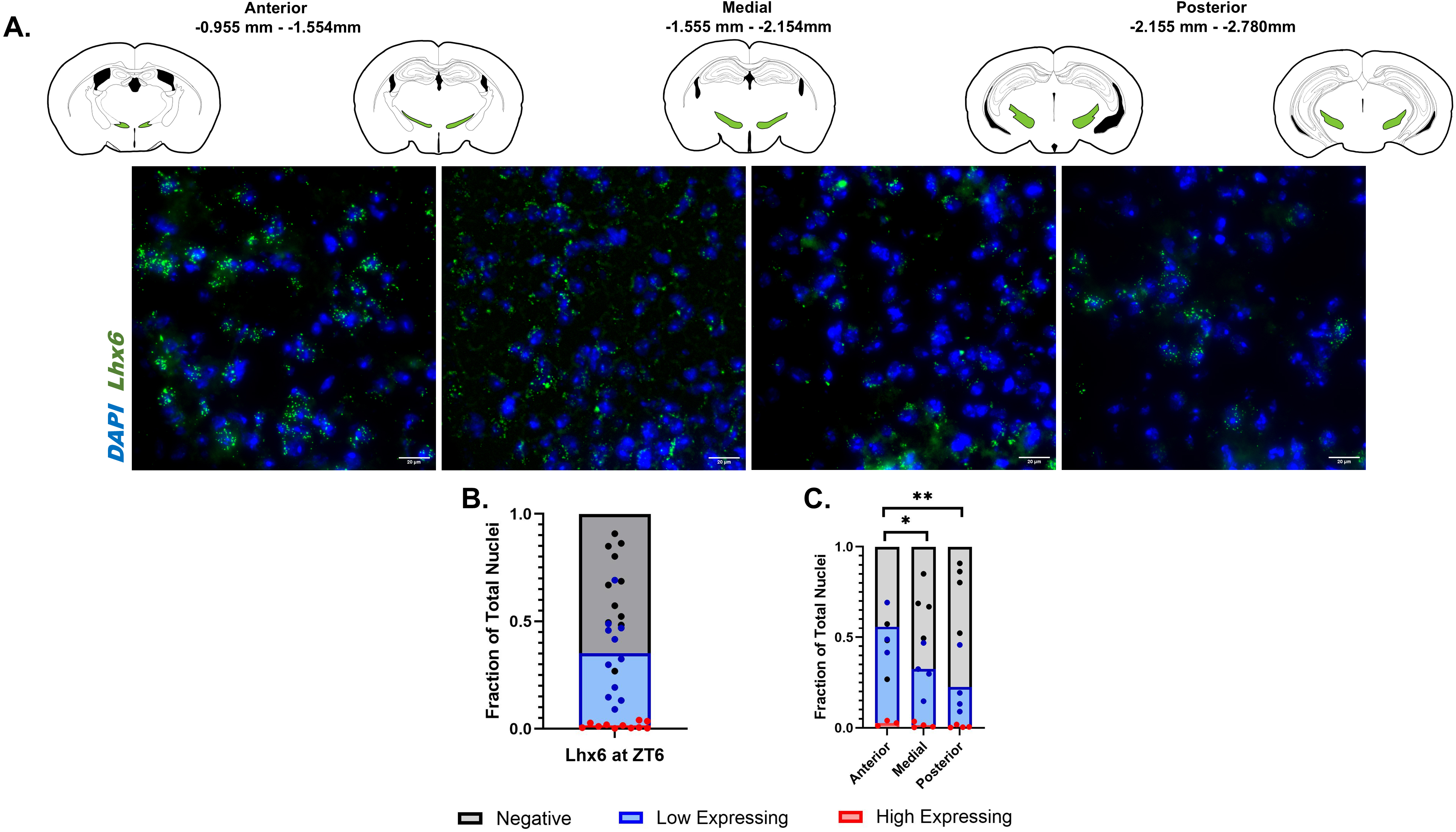
*Lhx6* expression decreases along the anterior-posterior axis of the zona incerta. **A.** Top row: Spatial diagram depicting the anterior and posterior limits of the zona incerta (green). Bottom row: *In situ* hybridization showing *Lhx6* (green) expressing cells (blue) decreasing along the anterior-posterior axis, covering a region spanning -0.955 mm to -2.780 mm Bregma. Scale bar=20 µm for all images**. B.** Bar graph depicting the fraction of *Lhx6*-positive cells at ZT6. *Lhx6-*positive cells compose 35.2% of total cells in the zona incerta (ZI), with high-expressing *Lhx6-*positive cells (red) comprising 1.5±1.3% of cells, low-expressing (blue) *Lhx6*-positive cells comprising 33.7±18.7% of cells, and the remaining 64.7±19.9% of cells do not express *Lhx6* (grey). **C.** Bar graph depicting the fraction of *Lhx6*-positive cells at ZT6 along the anterior-posterior axis. The anterior region contains the most *Lhx6*-positive expressing cells, with high-expressing cells comprising 2.6±1.5% of total cells and low-expressing cells comprising 53.2±14.3%. The medial region contains 1.5±1.4% high-expressing cells and 30.9±13.2% low-expressing cells. The posterior region contains the least *Lhx6*-positive cells, with high comprising 0.75±0.77% and low comprising 21.8±16.5%. There are significant differences in the numbers of low-expressing cells between the anterior and medial (*p* = 0.0187, alpha = 0.05) and the anterior and posterior (*p* = 0.0012, alpha=0.05) regions.

**Table 2:**
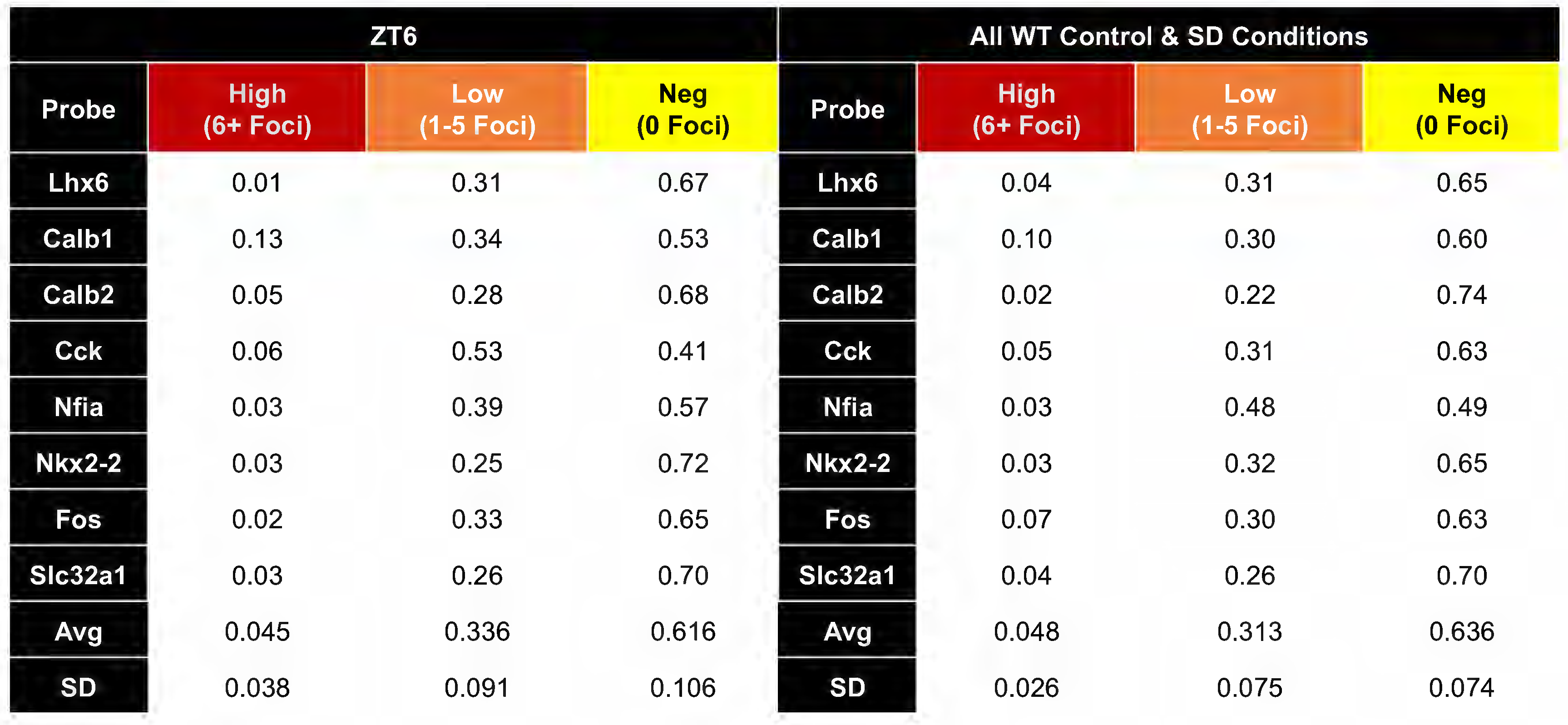
Puncta number distribution among probes tested. The table describes the composition of cell types amongst each marker at ZT6 (left) and amongst all experimental groups in wildtype mice (right). At ZT6, the majority of cells are negative for markers (61.6±10.6%), followed by low-expressing cells (33.6±9.1%). High expressing cells are the least abundant (4.5±3.8%). Similar trends exist across all experimental groups in wildtype animals (marker negative cells: 63.6±7.4%, low expressing cells: 31.3±7.5%, high expressing cells: 4.8±2.6%).

We performed the same analysis for the other markers. *Calb1* (11.9% high/33% low), *Cck* (4.8% high/42.5% low), and *Nfia* (3.9% high/38.8% low) labeled the largest fractions of ZI cells, although no probe labeled fewer than 25.8% of ZI cells, with *Nkx2-2* (3.7% high/22.5% low) showing the lowest overall prevalence (Fig. S3A, B). We observed only non-significant trends in the distribution of many of these markers along the anterior-posterior axis of the ZI, with *Fos* and *Nkx2-2* distribution paralleling the anterior enrichment seen for Lhx6, and *Calb1* exhibiting a medial bias (Fig. S3C).

### Responses of Lhx6-positive and Lhx6-negative ZI cells to sleep pressure

To profile activity changes under both naturally occurring and induced sleep pressure, we quantified *Fos* expression in both *Lhx6*-positive and *Lhx6*-negative ZI cells along the anterior-posterior axis of the ZI (Table 3; Supplemental Dataset 1). No significant changes in the total number of *Lhx6*-positive cells were detected between conditions (Fig. S4). Across the ZI, *Fos* levels were higher at high (ZT0) and mild-moderate (ZT6) sleep pressure than at low pressure (ZT12) (Fig. 2A,E) in both *Lhx6*-positive (Fig. 2B) and *Lhx6*-negative (Fig. 2C) populations, indicating that many ZI subtypes track naturally fluctuating sleep pressure.

**Figure 2:**
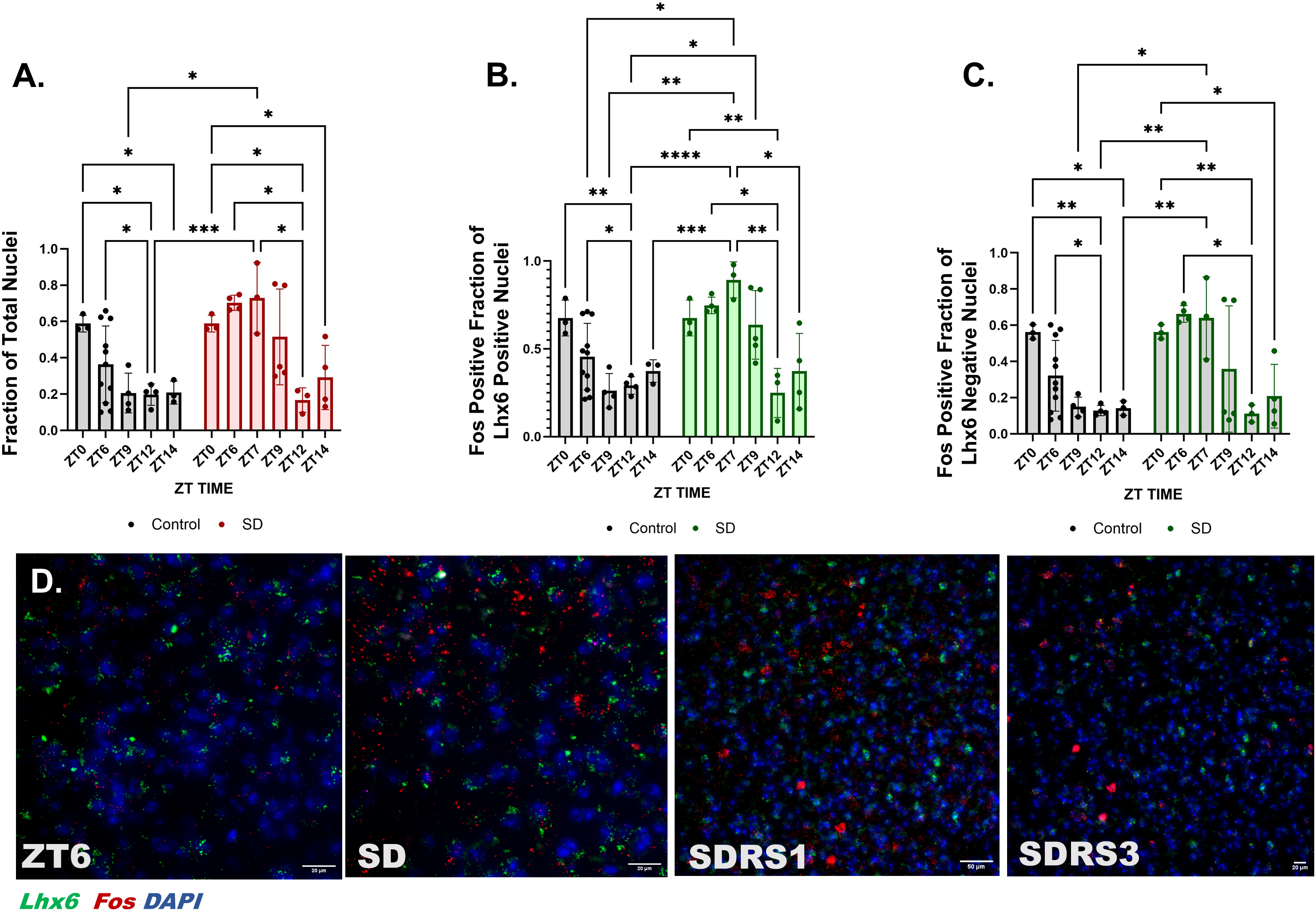
Induced sleep deprivation induces *Fos* expression in both *Lhx6*-positive and *Lhx6*-negative cells. **A.** Bar graph depicting the *Fos*-positive fraction of total cells in the undisturbed control and after sleep deprivation. Significant differences were found between: (alpha = 0.05) Control ZT0 v. Control ZT12 (*p* = 0.0117), Control ZT0 v. Control ZT14 (*p* = 0.0415), SD ZT0 v. SD ZT12 (SDRS6, *p* = 0.0117), SD ZT0 V. ZT14 SD (*p* = 0.0415), Control ZT6 v. Control ZT12 (*p* = 0.0274), SD ZT6 v. SD ZT12 (SDRS8, *p* = 0.0274), SD ZT7 (SDRS1) v. Control ZT9 (p = 0.0256), SD ZT7 (SDRS1) v. Control ZT12 (*p* = 0.0006), SD ZT7 (SDRS1) v. SD ZT12 (SDRS6) (*p* = 0.0393), SD ZT7 (SDRS1) v. Control ZT14 (*p* = 0.0022). **B.** Bar graph depicts the *Fos*-positive fraction of *Lhx6*-positive cells in control and sleep deprivation groups. Significant differences were found between: (alpha = 0.05) ZT0:Control v. ZT12:Control (*p* = 0.0068), ZT0:SD vs. ZT12:SD (*p* = 0.0068), ZT6:Control vs. ZT7:SD (*p* = 0.0221), ZT6:Control vs. ZT12:Control (*p* = 0.0261), ZT6:SD vs. ZT12:SD (*p* = 0.0261), ZT7:SD vs. ZT9:Control (*p* = 0.0033), ZT7:SD vs. ZT12:Control (*p* ≤ 0.0001), ZT7:SD vs. ZT12:SD (*p* = 0.0035), ZT7:SD vs. ZT14:Control (*p* = 0.0004), ZT7:SD vs. ZT14:SD (*p* = 0.0185), ZT9:SD vs. ZT12:Control (*p* = 0.0443). **C.** Bar graph depicting the *Fos*-positive fraction of *Lhx6*-negative cells in control and sleep deprivation groups. Significant differences were found: ZT0:Control vs. ZT12:Control (*p* = 0.0056), ZT0:Control vs. ZT14:Control (p = 0.0152), ZT0:SD vs. ZT12:SD (*p* = 0.0056), ZT0:SD vs. ZT14:SD (*p* = 0.0152), ZT6:Control vs. ZT12:Control (*p* = 0.0219), ZT6:SD vs. ZT12:SD (*p* = 0.0219), ZT7:SD vs. ZT9:Control (*p* = 0.0280), ZT7:SD vs. ZT12:Control (*p* = 0.0022), ZT7:SD vs. ZT14:Control (*p* = 0.0060). **D.** *In situ* hybridization showing *Lhx6* (green) and *Fos* (red) expression (DAPI. blue) in response to control (ZT6, scale bar=20 µm) and sleep deprivation conditions: sleep deprivation (SD, scale bar=20 µm), sleep deprivation with 1 hour recovery sleep (SDRS1, scale bar=50 µm), sleep deprivation with 3 hours recovery sleep (SDRS3, scale bar=20 µm). **Table 1** describes standard time and the corresponding zeitgeber time, and sleep deprivation conditions.

**Table 3:**
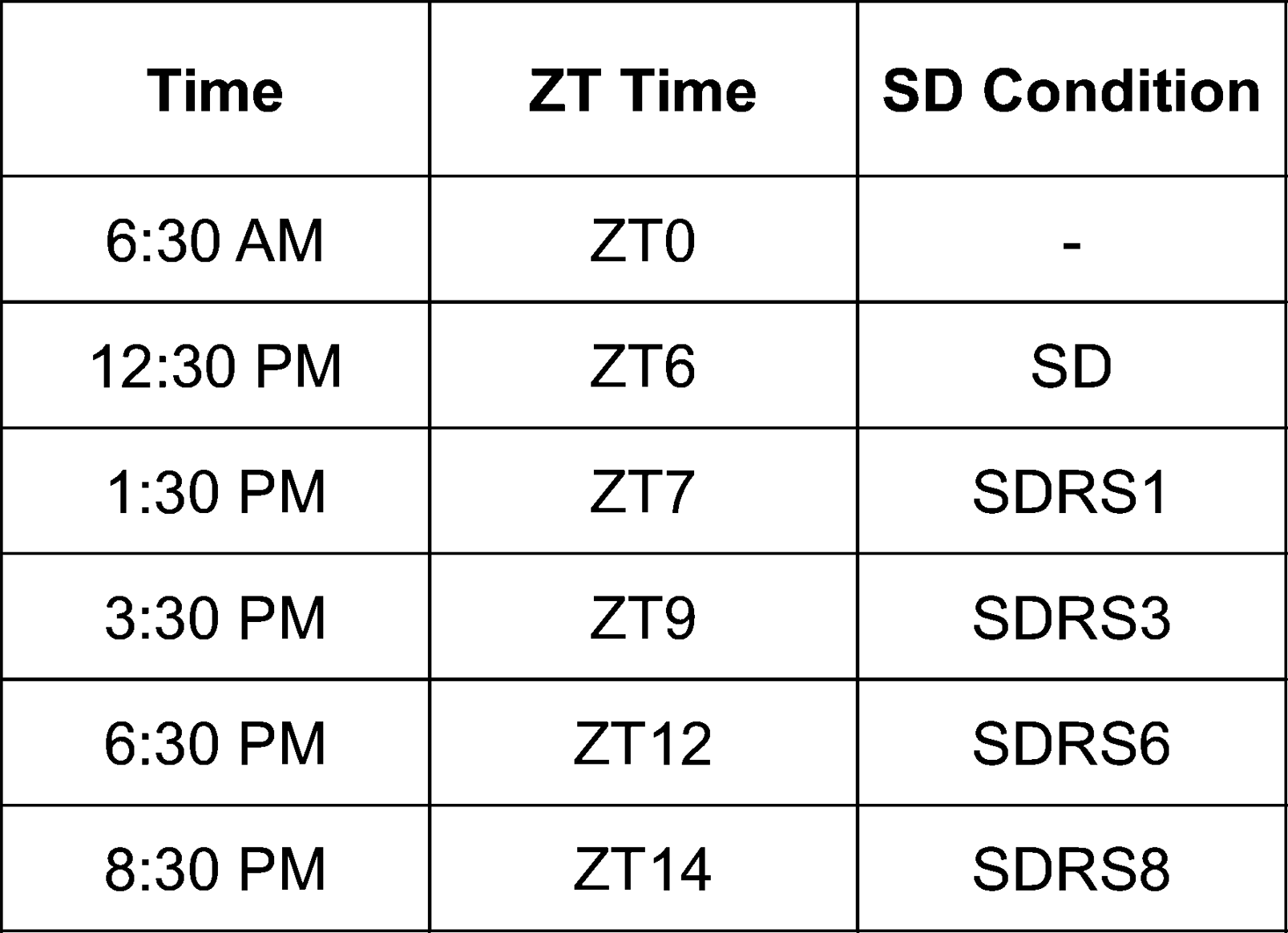
Samples examined by time and sleep deprivation conditions. The table lists the collection points during experimentation in standard time (left column), zeitberger time (middle column), and the corresponding sleep deprivation and recovery sleep conditions (right column).

To examine the effects of induced sleep pressure, mice were sleep deprived by continuous gentle brushing from ZT0 to ZT6 (Lemons, Saré, and Beebe Smith 2018; Liu et al. 2017a). One hour into recovery sleep (ZT7), *Fos* expression rose markedly in *Lhx6*-positive cells relative to ZT6 controls (Fig. 2B,E), but not in *Lhx6*-negative cells (Fig. 2C). After three hours of recovery (ZT9) *Fos* remained elevated, with a larger *Fos*-positive fraction in the *Lhx6*-positive cohort (Fig. 3A). By six hours of recovery (ZT12), *Fos* had declined in both groups (Fig. 2B,C). Notably, more *Lhx6*-positive neurons displayed high *Fos* intensity after sleep deprivation than their *Lhx6*-negative counterparts (Fig. 3C,F), demonstrating that Lhx6-positive cells mount a stronger and more persistent response as pressure dissipates (Fig. 3A-F).

**Figure 3:**
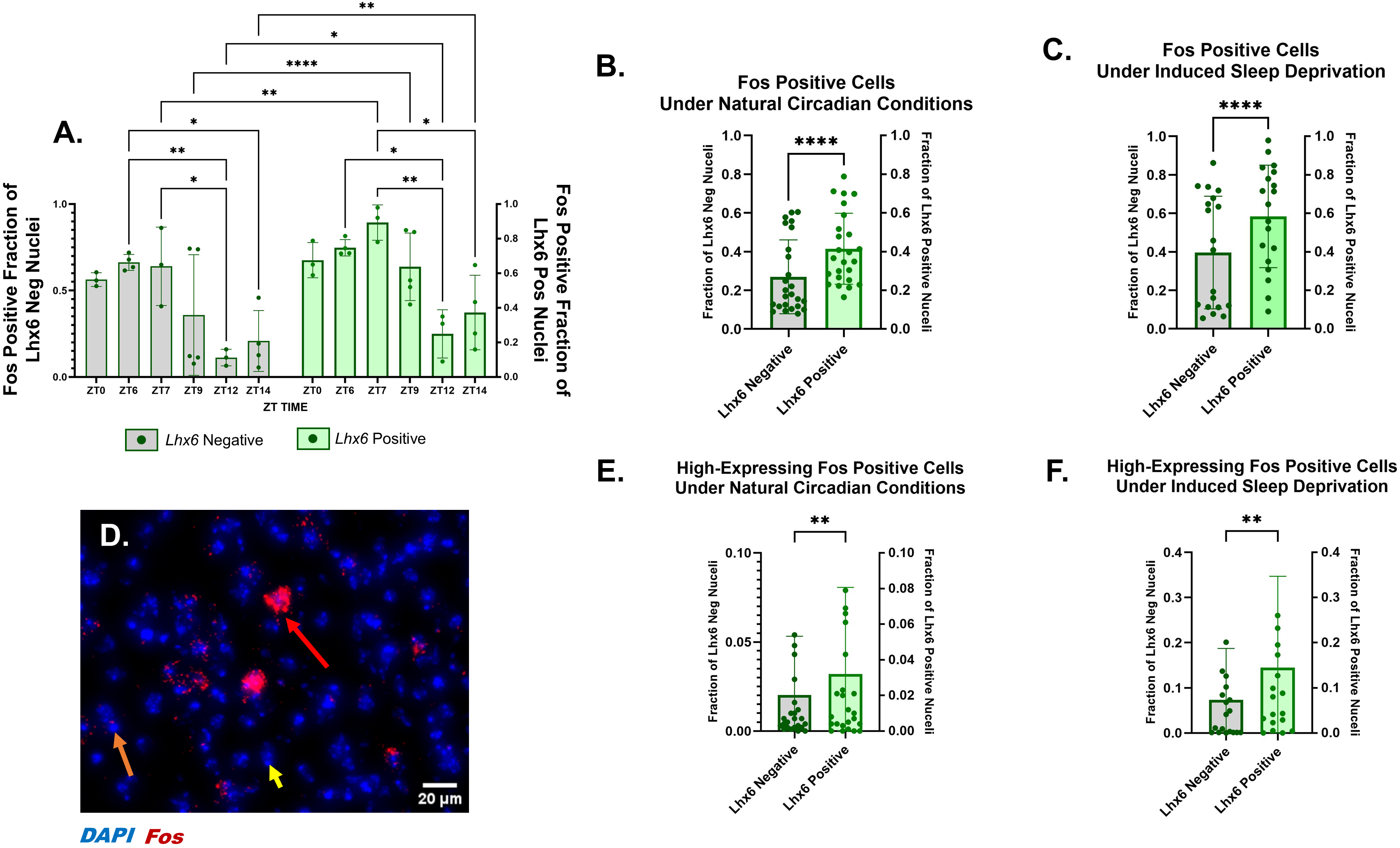
Sleep pressure-dependent *Fos* induction is stronger in *Lhx6*-positive than in *Lhx6*-negative cells. **A.** Bar graph depicts the *Fos*-positive fraction of *Lhx6*-negative cells (left, grey) and the Fos-positive fraction of Lhx6-positive cells (right, green) across ZT points. **B.** Bar graph depicts the *Fos*-positive fraction of *Lhx6*-negative cells (left, grey) and the *Fos*-positive fraction of Lhx6-positive cells (right, green) aggregated across all control timepoints under natural circadian conditions (*p* < 0.0001). **C.** Bar graph depicts the *Fos*-positive fraction of *Lhx6*-negative cells (left, grey) and the *Fos*-positive fraction of *Lhx6*-positive cells (right, green) aggregated across all control timepoints under induced sleep deprivation (p<0.0001). **D.** In situ hybridization illustrates examples of highly expressing Fos-positive cells (red arrow), low-expressing Fos cells (orange), and *Fos*-negative cells (orange). 20 µm scale bar. **E.** Bar graph depicts the high expressed *Fos*-positive fraction of Lhx6 negative cells (left, grey) and the Fos-positive fraction of *Lhx6*-positive cells (right, green) aggregated across all control timepoints under natural circadian conditions (*p* < 0.01). **F.** Bar graph depicts the high-expressing *Fos*-positive fraction of *Lhx6*-negative cells (left, grey) and the *Fos*-positive fraction of Lhx6-positive cells (right, green) aggregated across all control timepoints under induced sleep deprivation (*p* < 0.01).

Spatial analysis revealed that natural pressure rarely drove high *Fos* expression (> 5 puncta per cell) along the anterior–posterior axis, whereas early sleep deprivation induced numerous high-*Fos*/*Lhx6*-positive neurons, where *Fos* expression peaked after deprivation and one hour into recovery, then falling sharply after three hours of recovery sleep (Fig. S5). We conclude that induced sleep deprivation elicits much higher cellular levels of *Fos* in *Lhx6*-positive ZI neurons than does naturally arising sleep pressure changes.

### Responses of molecularly distinct subtypes of Lhx6-positive and Lhx6-negative ZI cells to sleep pressure

We next examine *Fos* induction in molecularly distinct subtypes of *Lhx6*-positive and *Lhx6*-negative ZI cells to sleep pressure (Table 3; Supplemental Dataset 1). As with *Lhx6* itself, none of the subtype markers showed any significant sleep pressure-dependent changes in gene expression (Fig. S6). *Cck* expression, however, did show time of day-dependent changes, with the highest expression observed at ZT0 and ZT6, and significantly reduced expression at later time points. However, the same pattern persisted during induced sleep pressure, indicating that while *Cck* expression may be regulated by circadian time and/or naturally occurring changes in sleep pressure, rather than pressure-dependent regulation.

All *Lhx6*-positive subtypes showed fewer *Fos*-positive cells under low sleep pressure (ZT12) than under high sleep pressure (ZT0) in unstimulated mice; all but the *Lhx6*-positive/*Calb2*-positive cells also differed between moderate (ZT6) and low sleep pressure (Fig. 4A-E).

**Figure 4:**
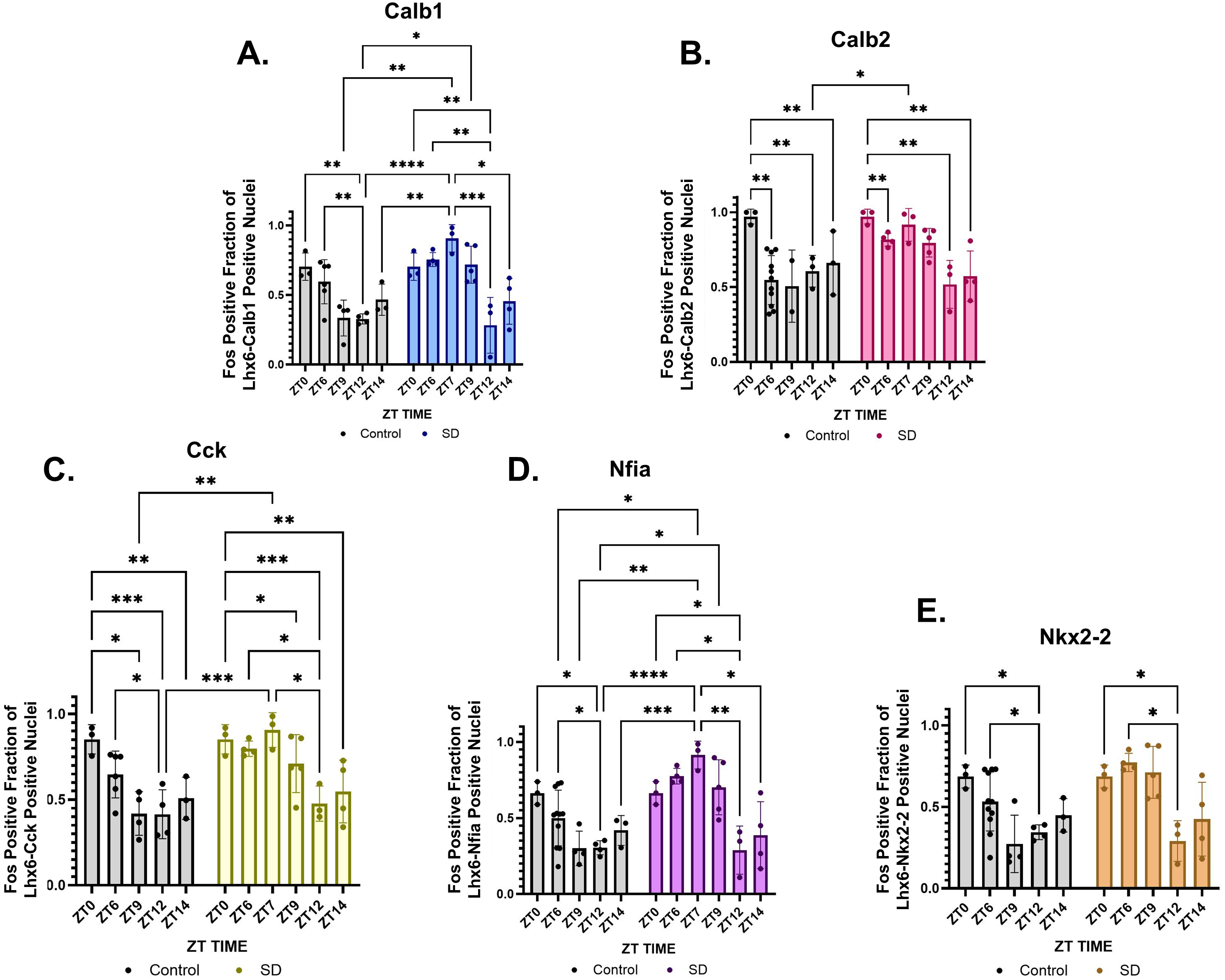
*Lhx6*-positive zona incerta cells expressing cell subtype-specific markers show similar changes in *Fos* induction in response to naturally-occurring and experimentally-induced changes in sleep pressure. Bar graphs depicting the *Fos*-positive fraction of marker-positive, *Lhx6*-positive cells under naturally occurring (grey) and induced sleep pressure (colored). **A.** *Calb1* (blue). Significant differences: ZT0:Control vs. ZT12:Control (*p* = 0.0020), ZT0:SD vs. ZT12:SD (*p* = 0.0020), ZT6:Control vs. ZT12:Control (*p* = 0.0012), ZT6:SD vs. ZT12:SD (*p* = 0.0012), ZT7:SD vs. ZT9:Control (*p* = 0.0086), ZT7:SD vs. ZT12:Control (*p* ≤ 0.0001), ZT7:SD vs. ZT12:SD (p = 0.0008), ZT7:SD vs. ZT14:Control (*p* = 0.0012), ZT7:SD vs. ZT14:SD (*p* = 0.0224), ZT9:SD vs. ZT12:Control (*p* = 0.0132). **B.** *Calb2* (red). Significant differences: ZT0:Control vs. ZT6:Control (*p* =0.0063), ZT0:Control vs. ZT12:Control (*p* = 0.0032), ZT0:Control vs. ZT14:Control (*p* =0.0078), ZT0:SD vs. ZT6:SD (*p* = 0.0063), ZT0:SD vs. ZT12:SD (*p* = 0.0032), ZT0:SD vs. ZT14:SD (*p* = 0.0078), ZT7:SD vs. ZT12:Control (*p* = 0.0431). **C.** *Cck* (yellow). Significant differences: ZT0:Control vs. ZT9:Control (*p* = 0.0171), ZT0:Control vs. ZT12:Control (*p* = 0.0003), ZT0:Control vs. ZT14:Control (*p* = 0.0045), ZT0:SD vs. ZT9:SD (p=0.0171), ZT0:SD vs. ZT12:SD (*p* = 0.0003), ZT0:SD vs. ZT14:SD (*p* = 0.0045), ZT6:Control vs. ZT12:Control (*p* = 0.0117), ZT6:SD vs. ZT12:SD (*p* = 0.0117), ZT7:SD vs. ZT9:Control (*p* = 0.0072), ZT7:SD vs. ZT12:Control (*p* = 0.0002), ZT7:SD vs. ZT12:SD (p*p* = 0.0106), ZT7:SD vs. ZT14:Control (*p* = 0.0023) **D**. *Nfia* (purple). Significant differences: ZT0:Control vs. ZT12:Control (p=0.0181), ZT0:SD vs. ZT12:SD (*p* = 0.0181), ZT6:Control vs. ZT7:SD (*p* = 0.0328), ZT6:Control vs. ZT12:Control (*p* = 0.0137), ZT6:SD vs. ZT12:SD (*p* = 0.0137), ZT7:SD vs. ZT9:Control (*p* = 0.0069), ZT7:SD vs. ZT12:Control (*p* < 0.0001), ZT7:SD vs. ZT12:SD (0.0034), ZT7:SD vs. ZT14:Control (*p* = 0.0004), ZT7:SD vs. ZT14:SD (*p* = 0.0199) **E.** *Nkx2-2* (orange) ZT0:Control vs. ZT12:Control (*p* = 0.0215), ZT0:SD vs. ZT12:SD (*p* = 0.0215), ZT6:Control vs. ZT12:Control (*p* = 0.0169), ZT6:SD vs. ZT12:SD (p=0.0169).

Following experimentally-induced sleep deprivation, significant decreases in *Fos* were observed between samples at ZT6 (end of SD treatment) and/or ZT7 (SD with 1 hour of recovery sleep) relative to ZT12 (SD with 6 hours of recovery sleep). This decreased Fos expression was observed in every *Lhx6*-positive subtype except for *Lhx6*-positive/*Calb2*-positive cells (Fig. 4A-E). *Lhx6*-negative neurons bearing these markers largely mirrored these kinetics (Fig. S7A-E), with two exceptions: *Cck*- and *Calb2*-expressing *Lhx6*-negative cells followed a circadian-like pattern, indicating weaker responsiveness to induced deprivation (Fig. S7B-C).

### *Nkx2-2* is essential for the development and function of Lhx6-positive ZI neurons

These data indicate that *Lhx6-*positive/*Nkx2-2*-positive cells may be particularly sensitive to sleep pressure. Previous studies from our group have also suggested that *Nkx2-2* may be important for the specification and/or differentiation of a subset of hypothalamic *Lhx6*-positive neurons, including those in the ZI (Kim et al. 2021b). To test this directly, we generated *Lhx6-Cre;Nkx2-2^lox/lox^* mice, thereby selectively inactivating *Nkx2-2* in *Lhx6*-expressing cells (Fragkouli et al. 2009; Mastracci, Lin, and Sussel 2013; Gross et al. 2016) (Fig. 5A). Mouse litters contained the expected Mendelian ratios of *Lhx6-Cre;Nkx2-2^lox/l+^* and *Lhx6-Cre;Nkx2-2^lox/lox^*offspring (Fig. S10).

**Figure 5:**
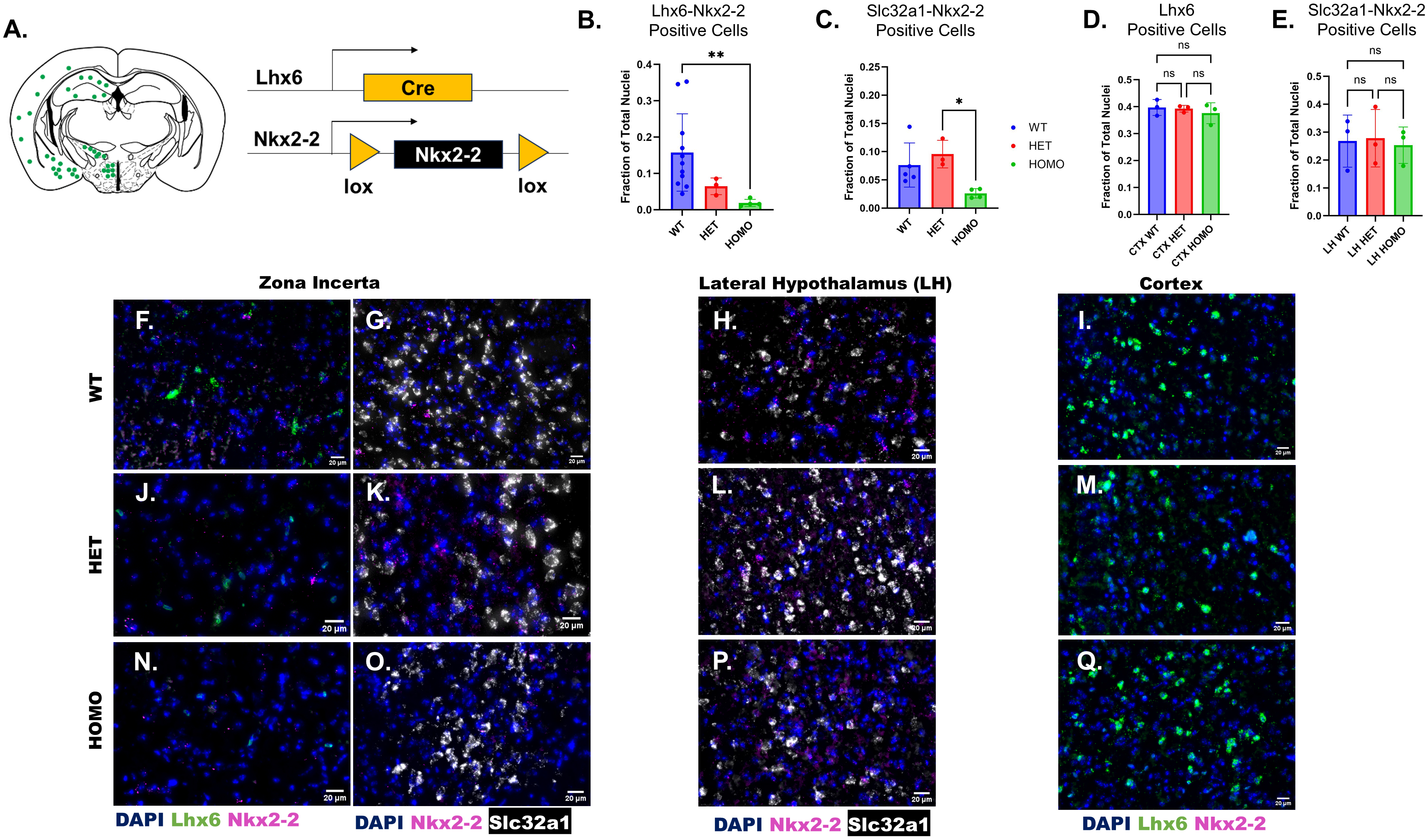
*Lhx6-Cre;Nkx2-2^lox/lox^* mice show reduced expression of both *Lhx6* and *Nkx2-2* in zona incerta. **A.** Diagram depicting the expression of *Lhx6* (green) in the coronally sliced mouse brain (left). Diagram depicting Lhx6-Cre-mediated deletion of *Nkx2-2* (right). **B.** Bar graph depicting the fraction of *Lhx6-Nkx2-2* positive cells of total cells amongst experimental groups: Wildtype (WT, blue), *Lhx6-Cre;Nkx2-2^lox/+^* (HET, red), and *Lhx6-Cre;Nkx2-2^lox/lox^*(HOMO, green). Significant differences: WT vs. HOMO (p = 0.0038). **C.** Bar graph depicting the fraction of *Slc32a1*/*Nkx2-2* positive cells of total cells amongst experimental groups: Wildtype (WT, blue), *Lhx6-Cre;Nkx2- 2^lox/+^* (HET, red), and *Lhx6-Cre;Nkx2-2^lox/lox^*(HOMO, green). Significant differences: HET vs. HOMO (p = 0.0194). **D.** Bar graph depicting the fraction of *Lhx6*-positive cells of total cells in the cortex (CTX) amongst experimental groups: Wildtype (WT, blue), *Lhx6-Cre;Nkx2-2^lox/+^* (HET, red), and *Lhx6-Cre;Nkx2-2^lox/lox^*(HOMO, green). **E.** Bar graph depicting the fraction of *Slc32a1*/Nkx2-2 positive cells of total cells in the lateral hypothalamus (LH) amongst experimental groups: Wildtype (WT, blue), *Lhx6-Cre;Nkx2-2^lox/+^* (HET, red), and *Lhx6-Cre;Nkx2- 2^lox/lox^*(HOMO, green). **F-I.** *In situ* hybridization depicting the expression of *Lhx6* (green), *Nkx2-2* (purple), *Slc32a1* (white) in cells (DAPI, blue) in the Zona Incerta (ZI, F-G), lateral hypothalamus (LH, H), and the Cortex (CTX, I) in wildtype (WT) mice. **J-M.** *In situ* hybridization depicting the expression of *Lhx6* (green), *Nkx2-2* (purple), Slc32a1 (white) in cells (DAPI, blue) in the ZI (J-K), LH (L), and cerebral cortex (M) in *Lhx6-Cre;Nkx2-2^lox/+^* (HET) mice. **N-Q.** *In situ* hybridization depicting the expression of *Lhx6* (green), *Nkx2-2* (purple), *Slc32a1* (white) in cells (DAPI, blue) in the ZI (J-K), LH (L), and cortex (M) in *Lhx6-Cre;Nkx2-2^lox/lox^* (HOMO) mice. Scale bar=20 µm for all images.

Mutant animals were indistinguishable from controls in appearance, body weight, and general locomotor activity. In the ZI, however, *Lhx6-Cre;Nkx2-2^lox/lox^* mice showed significant reductions in both *Lhx6*-positive/*Nkx2-2*-positive and *Slc32a1*-positive/*Nkx2-2*-positive GABAergic neurons, indicating the efficiency of the intersectional loss of function mutants generated here (Fig. 5B, C). Specificity was confirmed by the finding of unchanged numbers of both *Lhx6*-positive cortical interneurons, which never express *Nkx2-2*, and *Slc32a1*-positive/*Nkx2-2*-positive GABAergic neurons in the lateral hypothalamus, where *Lhx6-Cre* is inactive (Fig. 5D).

A non-significant trend towards reduced numbers of *Lhx6*-positive/*Nkx2-2*-positive cells was also observed in heterozygous *Lhx6-Cre;Nkx2-2^lox/+^* mice, indicating a possible dose-dependent requirement for *Nkx2-2* in the development of *Lhx6*-positive ZI neurons (Fig. 5B, C). Loss of both *Lhx6*-positive and *Nkx2-2*-positive cells was evident across all positions along the anterior-posterior axis of the ZI (Fig. S11).

Loss of *Nkx2-2* also disrupted expression of other subtype markers of Lhx6-positive ZI neurons (Fig. 6). Relative to wild-type animals, both heterozygous *Lhx6-Cre;Nkx2-2^lox/+^* and homozygous *Lhx6-Cre;Nkx2-2^lox/lox^*mutants displayed significant reductions in *Calb1-*, *Calb2-*, and *Cck-*positive Lhx6 neurons. in. We also observe a trend towards reduced expression of all probes in the medial ZI, and a corresponding relative increase in both the anterior and, in particular, the posterior ZI. In heterozygous *Lhx6-Cre;Nkx2-2^lox/+^* mice, a similar redistribution was observed for *Lhx6*. Although the total number of *Slc32a1*-positive cells in the ZI was unchanged across all genotypes examined, the relative number of cells showing strong (>5 puncta/cell) *Slc32a1* expression was reduced in homozygous *Lhx6-Cre;Nkx2-2^lox/lox^* mutants, relative to *Lhx6-Cre;Nkx2-2^lox/+^* heterozygotes.

**Figure 6:**
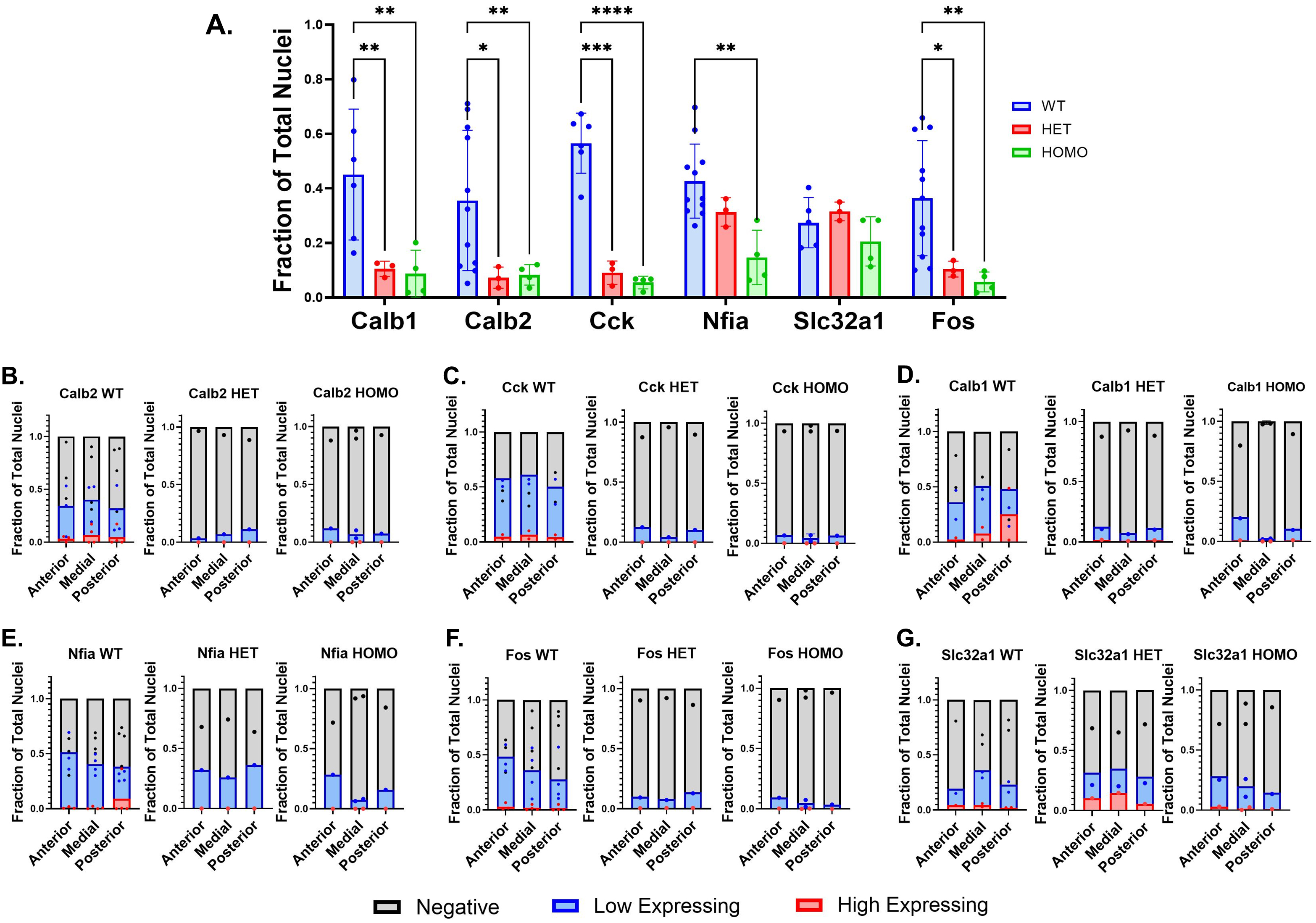
*Lhx6-Cre;Nkx2-2^lox/lox^* mice show reduced expression of cell subtype-specific markers in zona incerta. **A.** Bar graph depicting the marker (from left to right: Calb1, Calb2, Cck, Nfia, Slc32a1, Fos) positive fraction of total cells across experimental groups: Wildtype (WT, blue), *Lhx6-Cre;Nkx2- 2^lox/+^* (HET, red), *Lhx6-Cre;Nkx2-2^lox/lox^* (HOMO, green). Significant differences were found between wildtype and het in the expression of Calb1 (*p* = 0.0065), Calb2 (*p* = 0.0175), Cck(*p* = 0.0001), and Fos (*p* = 0.0320); as well as between wildtype and homo in the expression of Calb1(p=0.0015), Calb2(*p* = 0.0098), Cck(*p* = <0.0001), Nfia(*p* = 0.0078), and Fos(*p* = 0.0032). **B-G.** Bar graphs depicting the fraction of marker-positive cells (high expressing, red; low expressing, blue) of total cells, along the anterior to posterior axis, across experimental groups. Right panel: Wildtype (WT), Middle panel: *Lhx6-Cre;Nkx2-2^lox/+^* (HET), left panel: *Lhx6-Cre;Nkx2-2^lox/lox^* (HOMO). No statistical analysis conducted due to HET and HOMO samples containing N < 2 per location.

We next used Xenium-based spatial transcriptomic analysis to more comprehensively analyze gene expression in wildtype mice using a 347-gene panel, which consisted of the Xenium Mouse Brain probeset (Ma et al. 2024) and 100 additional probes selected for enriched expression in hypothalamic cell types (Kim et al. 2020). After segmenting cells from the ZI and adjacent tissues and performing UMAP analysis on their expression profiles, we resolved discrete clusters of glutamatergic and GABAergic neurons, and non-neuronal cell types such as astrocytes, oligodendrocyte precursor cells, mature oligodendrocytes, endothelial cells, and microglia (Fig. 7C). Subclustering the GABAergic *Lhx6*-expressing neuronal population identified a subset corresponding to *Lhx6*-positive ZI neurons (Fig. 7E-G). This analysis confirmed the anterior-posterior gradient in the relative number of *Lhx6*-expressing cells in wildtype animals observed using HiPlex analysis (Fig. 7H). We identified additional genes enriched in *Nkx2-2*-positive and *Nkx2-2*-negative cell types, and further subclusters within these groups (Fig. S12). *Nkx2-2*-positive cells comprised four clusters (clusters 0-3), and expressed *Lypd6, Nkx2-1, Foxp2, Ly6a, Adgrl4, Paqr5, Penk, and Dlx1*. In contrast, *Nkx2-2*-negative cell types (clusters 4 and 5) lack expression of these markers. Instead, *Nkx2-2*-negative cell types have more intense and abundant expression of *Col6a1* and *Tmem132d*, and more abundant expression of *Calb2*.

**Figure 7:**
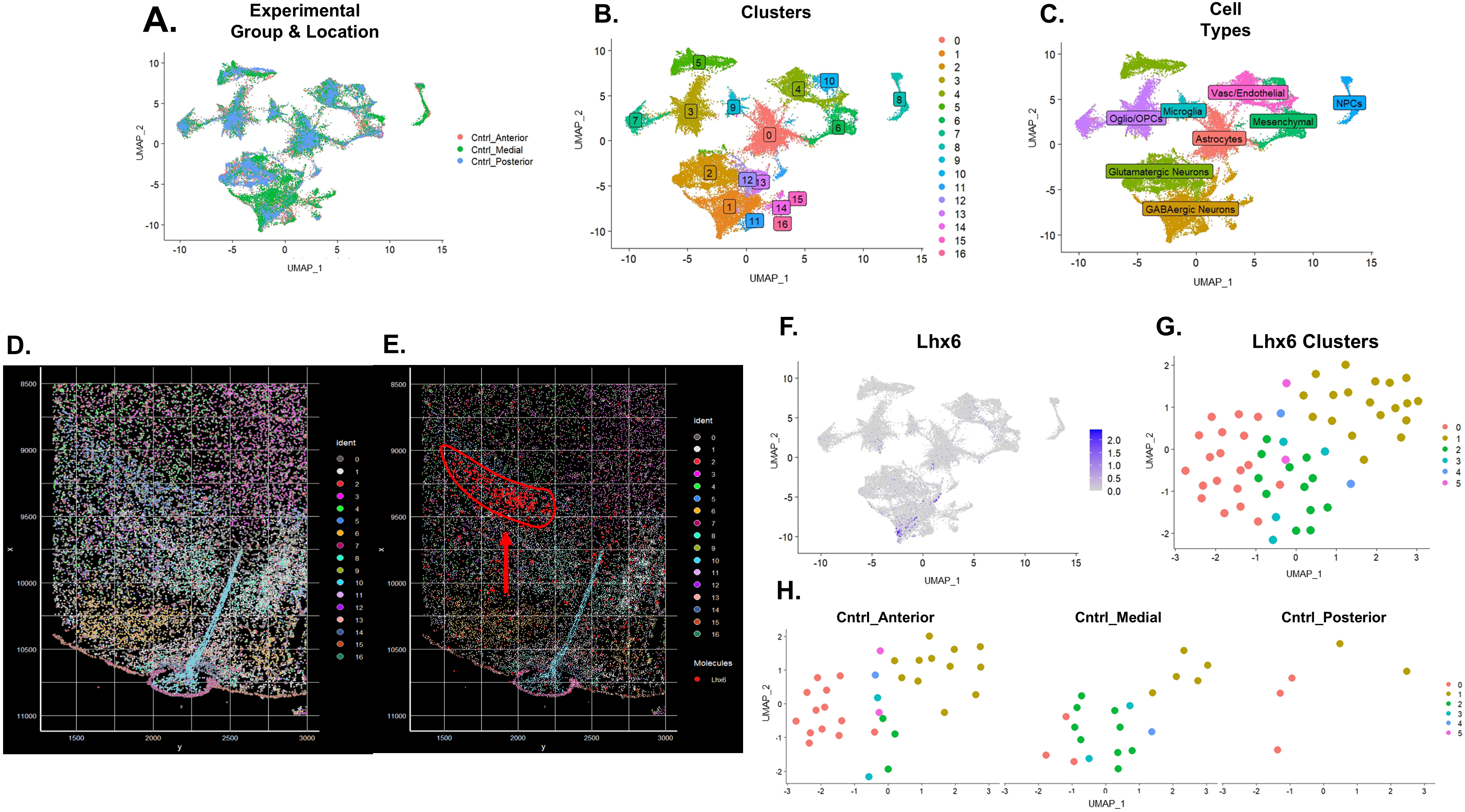
Xenium-based analysis of *Lhx6*-positive ZI cells. **A.** UMAP depicts the clustering of wild-type anterior, medial, and posterior samples. **B.** UMAP identifies 17 clusters in wildtype anterior, medial, and posterior samples. **C.** UMAP depicts major cell types. **D**. Visualization of Seurat-identified clusters in the spatial context of the hypothalamus. **E.** Visualization of Seurat identified clusters in the spatial context of the hypothalamus with *Lhx6*-expressing ZI neurons highlighted (red with red arrow). **F.** UMAP identifying the GABAergic *Lhx6* population. **G**. UMAP depicts subclusters of GABAergic *Lhx6-*positive cells after additional filtration of cells expressing astrocyte, glial, and glutamatergic markers. **H.** UMAP depicts subclusters of GABAergic *Lhx6* neurons distributed along the anterior-posterior axis.

We next tested whether the reduction in the number of *Lhx6*-positive cells was reflected in defective activation of the remaining ZI cells to elevate sleep pressure. In both heterozygous *Lhx6-Cre;Nkx2-2^lox/+^*and homozygous *Lhx6-Cre;Nkx2-2^lox/lox^* mutants, we observed dramatic reductions in the number of total *Fos*-positive neurons under conditions of both moderate (ZT6), moderately high (SD+3 hours recovery sleep) and high (ZT6+6 hours of SD) sleep pressure (Fig.8). The few remaining *Lhx6*-positive/*Nkx2-2*-positive ZI cells likewise showed reduced levels of overall activation, and no clear changes in activity in response to altered sleep pressure. High levels of *Fos* expression were likewise not observed in cells at any point along the anterior-posterior axis of the ZI (Fig. S13).

**Figure 8:**
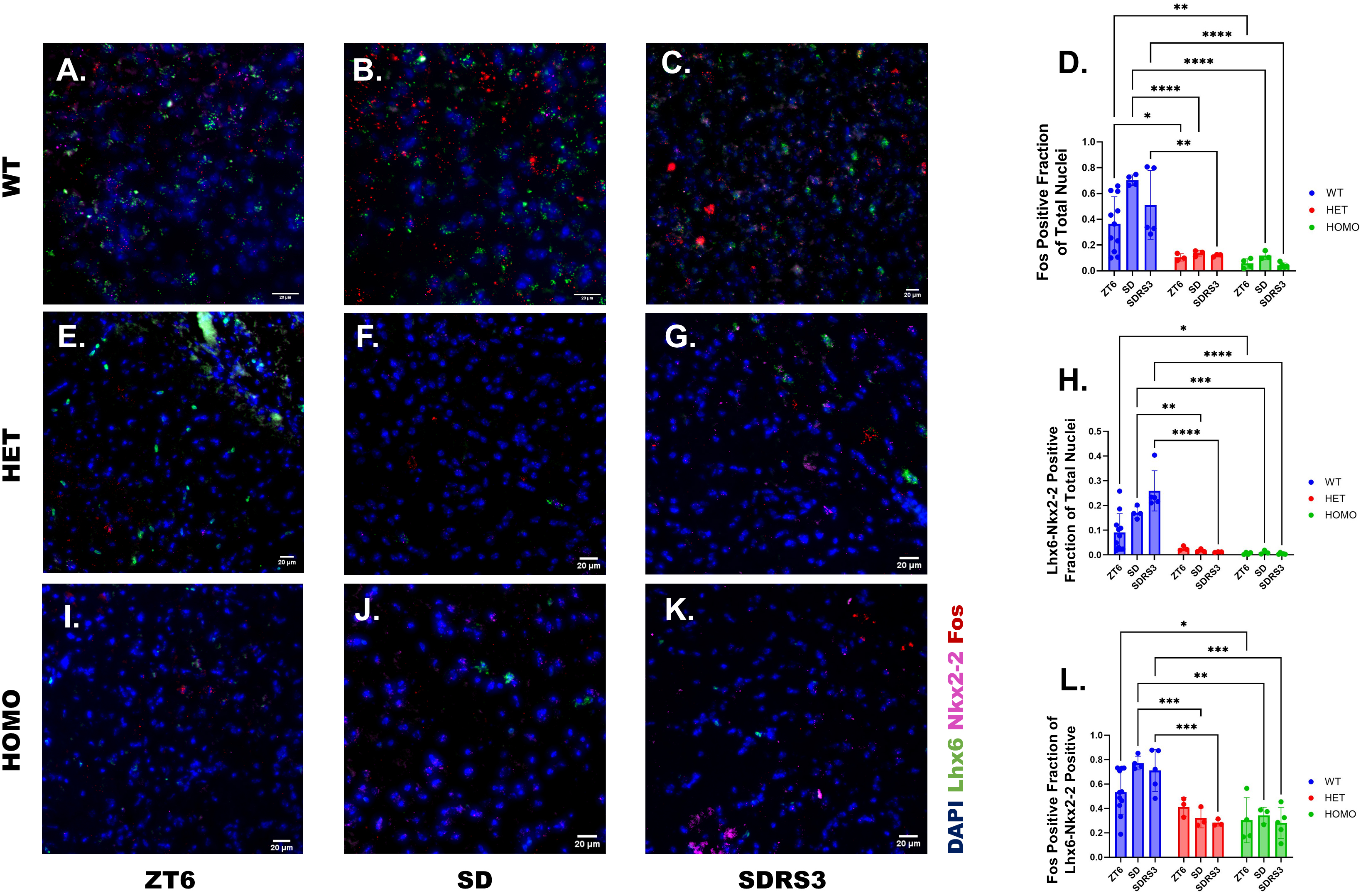
*Lhx6-Cre;Nkx2-2^lox/lox^* mice fail to induce *Fos* in response to experimentally-induced sleep deprivation. **A-C.** *In situ* hybridization depicts *Fos* (red) and *Lhx6* (green) expression in wildtype (WT) at ZT6 (A), after sleep deprivation (SD, B), and after sleep deprivation with 3 hours of recovery sleep (SDRS3, C). **D.** Bar graph depicts the *Fos*-positive fraction of total cells in wildtype (WT, blue), *Lhx6-Cre;Nkx2-2^lox/+^* (HET, red), and *Lhx6-Cre;Nkx2-2^lox/lox^*(HOMO, green) at ZT6, SD, and SDRS3. Significant differences were found between the wildtype and HET at ZT6(p=0.0354), SD(p<0.0001), and SDRS3(p=0.0037); as well as the wildtype and HOMO at ZT6(p=0.0045), SD(p<0.0001), and SDRS3(p<0.0001). **E-G.** *In situ* hybridization depicts *Fos* (red) and *Lhx6* (green) expression in *Lhx6-Cre;Nkx2-2^lox/+^* (HET) mice at ZT6 (E), after sleep deprivation (SD, F), and after sleep deprivation with 3 hours of recovery sleep (SDRS3, G). **H.** Bar graph depicts the *Lhx6-Nkx2-2* positive fraction of total cells in wildtype (WT, blue), *Lhx6-Cre;Nkx2-2^lox/+^* (HET, red), and *Lhx6-Cre;Nkx2-2^lox/lox^*(HOMO, green) at ZT6, SD, and SDRS3. Significant differences were found between wildtype and HET at SD (p=0.0015) and SDRS3 (p<0.0001); as well as the wildtype and HOMO at ZT6(0.0235), SD(p=0.0009), and SDRS3(p<0.0001). **I-K.** *In situ* hybridization depicts *Fos* (red) and *Lhx6* (green) expression in *Lhx6-Cre;Nkx2-2^lox/lox^* (HOMO) mice at ZT6 (I), after sleep deprivation (SD, J), and after sleep deprivation with 3 hours of recovery sleep (SDRS3, K). **L.** Bar graph depicts the *Fos* positive fraction of *Lhx6-Nkx2-2* positive cells in wildtype (WT, blue), *Lhx6-Cre;Nkx2-2^lox/+^* (HET, red), and *Lhx6-Cre;Nkx2-2^lox/lox^*(HOMO, green) at ZT6, SD, and SDRS3. Significant differences were found between the wildtype and HET at SD(p=0.0007) and SDRS3(p=0.0008); as well as the wildtype and HOMO at ZT6(p=0.0259), SD(p=0.0012), and SDRS3(p=0.0001).

Finally, we examined sleep patterns in control *Lhx6-Cre* mice and heterozygous *Lhx6-Cre;Nkx2-2^lox/+^* and homozygous *Lhx6-Cre;Nkx2-2^lox/lox^*mutants using the Piezo system (Yaghouby et al. 2016) (Fig. 9). Relative to control animals, heterozygous *Lhx6-Cre;Nkx2-2^lox/+^* mice showed increased sleep time during the day, but no change in total nighttime sleep, or either daytime or nighttime sleep bout length. In contrast, homozygous *Lhx6-Cre;Nkx2-2^lox/lox^* showed significantly increased sleep time and sleep bout length during both day and night relative to both heterozygotes and, except for nighttime sleep bout length, to controls.

**Figure 9:**
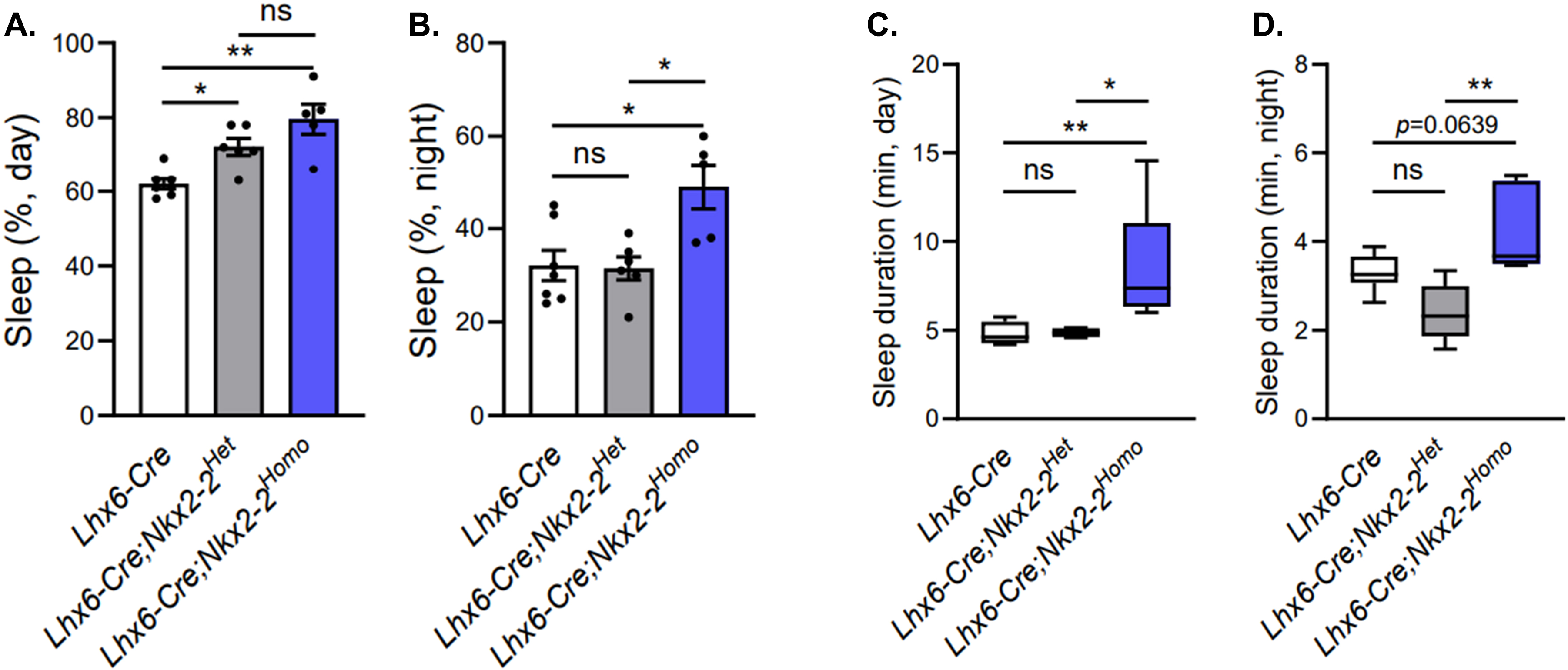
Total sleep is increased in *Lhx6-Cre;Nkx2-2^lox/+^* and *Lhx6-Cre;Nkx2-2^lox/lox^* mice. A Bar graph depicts the percent of time spent asleep during the day in *Lhx6-Cre* (white), *Lhx6-Cre;Nkx2-2^lox/+^* (grey), and *Lhx6-Cre;Nkx2-2^lox/lox^*mice (blue). Significant differences found between *Lhx6-Cre* and *Lhx6-Cre;Nkx2-2^lox/+^* mice (p≤0.05)as well as between *Lhx6-Cre* and *Lhx6-Cre;Nkx2-2^lox/lox^* mice (p≤ 0.01). B Bar graph depicts the percent of time spent asleep at night in *Lhx6-Cre* (white), *Lhx6-Cre;Nkx2-2^lox/+^*(grey), and *Lhx6-Cre;Nkx2-2^lox/lox^* mice (blue). Significant differences were found between *Lhx6-Cre* and *Lhx6-Cre;Nkx2-2^lox/lox^* mice (p≤0.05), as well as between *Lhx6-Cre;Nkx2-2^lox^*^/+^ and *Lhx6-Cre;Nkx2-2^lox/lox^* mice (p≤0.05). C Box and whisker plot depicts the duration of sleep bouts during the day in *Lhx6-Cre* (white), *Lhx6-Cre;Nkx2-2^lox/+^* (grey), and *Lhx6-Cre;Nkx2-2^lox/lox^* mice (blue). Significant differences were found between *Lhx6-Cre* and *Lhx6-Cre;Nkx2-2^lox/lox^*mice (p≤0.01), as well as between *Lhx6-Cre;Nkx2-2^lox/+^* and *Lhx6-Cre;Nkx2-2^lox/lox^* mice (p≤0.05). C Box and whisker plot depicts the duration of sleep bouts at night in *Lhx6-Cre* (white), *Lhx6-Cre;Nkx2-2^lox/+^* (grey), and *Lhx6-Cre;Nkx2-2^lox/lox^* mice (blue). A significant difference was found between *Lhx6-Cre;Nkx2-2^lox/+^* and Lhx6-Cre;Nkx2-2^lox/lox^ mice.

## Discussion

The ZI was first studied for its role in sensorimotor integration (Deschênes et al. 2005; Chometton, Barbier, and Risold 2021; Hormigo et al. 2023; Schäfer and Hoebeek 2018). More recently, it has emerged as a hub for regulating internal states and innate behaviours, including feeding, thirst, and anxiety and aggression-related behaviors (de Git et al. 2021; Zhao et al. 2019; Li, Rizzi, and Tan 2021; Walsh and Grossman 1976; Chou et al. 2018) (de Git et al. 2021; Zhao et al. 2019; Li, Rizzi, and Tan 2021; Walsh and Grossman 1976; Chou et al. 2018).

Importantly, the ZI also contributes to sleep-wake regulation by detecting and signaling sleep pressure (Liu et al. 2017b; Vidal-Ortiz, Blanco-Centurion, and Shiromani 2024; Lee et al. 2025; Blanco-Centurion et al. 2021). *Lhx6*-positive ZI neurons are critical for these functions: developmental disruption of *Lhx6* function causes apoptotic death, reduces their numbers, and decreases overall sleep (Kim et al. 2021a; Liu et al. 2017a). Glutamatergic neurons of the thalamic reuniens nucleus, which mediate the effects of induced sleep pressure, directly activate anterior Lhx6-positive ZI neurons (Lee et al. 2025). However, activation patterns within *Lhx6*-positive neurons are heterogeneous, and responses of *Lhx6*-negative ZI neurons to sleep pressure remain poorly characterized.

### Sleep pressure-dependent activation of *Lhx6*-positive and *Lhx6*-negative ZI neurons

Using multiplex smFISH, we quantified *Fos* induction in *Lhx6*-positive and *Lhx6*-negative ZI neurons across molecular subtypes and examined developmental and behavioral effects of selectively disrupting *Nkx2-2* in *Lhx6*-positive neurons. Consistent with prior reports (Liu et al. 2017a), *Lhx6*-positive neurons robustly induced *Fos* following naturally occurring and experimentally-induced sleep pressure, and sustained elevated expression for at least 3 hours into recovery sleep before gradually declining. Induced sleep pressure initially triggered higher *Fos* levels than naturally-regulated sleep pressure changes, though this difference disappeared after several hours of recovery sleep. *Lhx6*-negative neurons showed similar but weaker *Fos* responses, with fewer *Fos*-positive cells overall and lower cellular expression levels. Both *Lhx6*-positive and -negative anterior ZI neurons exhibited higher *Fos* induction than posterior ZI neurons, paralleling the pattern of reuniens inputs (Lee et al. 2025). We conclude that while *Lhx6*-positive neurons show the strongest responses to elevated sleep pressure, multiple ZI subtypes participate in sleep pressure signaling.

### Molecular diversity of sleep pressure-responsive ZI neuronal subtypes

ZI neurons are highly heterogeneous (M. Zhang et al. 2023), yet activity-dependent differences among subtypes remain largely unexplored. Using a panel of molecular markers, we observed no sleep pressure-dependent changes in their expression, though *Cck* levels rose at the start of the circadian day. *Nkx2-2* and *Nfia* were enriched anteriorly, while *Calb1*, *Calb2*, and *Cck* were more medial/posterior. *Fos* induction across these molecularly defined subtypes largely paralleled patterns in *Lhx6*-positive neurons, with *Lhx6*-negative counterparts showing reduced numbers of *Fos*-positive cells and lower expression levels. These findings confirm that sleep pressure broadly activates diverse ZI cell types, with *Lhx6*-positive neurons showing the strongest responses.

### Nkx2-2 and developmental specification of Lhx6-positive ZI neurons

To define Nkx2-2’s role in ZI development, we disrupted it selectively in *Lhx6*-positive neurons and analyzed molecular and behavioral consequences using both HiPlex smFISH and Xenium spatial transcriptomics. Wildtype ZI neurons segregated into *Nkx2-2*-positive and -negative groups with distinct molecular markers. *Nkx2-2* loss decreased expression of markers for *Nkx2-2*-positive neurons and increased those for *Nkx2-2*-negative neurons, with residual *Nkx2-2*-positive populations displaced anteriorly and posteriorly. *Calb1*, *Calb2*, and *Cck* expression was reduced in both heterozygous and homozygous mutants, while overall *Lhx6*-positive neuron numbers declined significantly in homozygotes. These results imply that *Nkx2-2* acts during development to specify both *Lhx6*-positive and -negative ZI neurons.

### Consequences of Nkx2-2 loss for sleep pressure signaling and sleep behavior

Developmental Nkx2-2 loss of function produced profound effects on sleep pressure signaling and behavior. *Fos* induction in *Lhx6*-positive neurons was markedly reduced under all conditions, especially in homozygous mutants. Paradoxically, these animals exhibited increased sleep time and bout length, with homozygotes showing stronger effects. Thus, Nkx2-2 disruption compromises ZI signaling of sleep pressure and alters sleep architecture, with effects opposite to those observed following complete Lhx6 loss.

### Coordinated sleep pressure coding in the ZI

These results underscore a central role for *Lhx6*-positive ZI neurons in signaling levels of both naturally occurring and induced sleep pressure, but also show that many *Lhx6*-negative ZI cells show broadly similar responses to elevated sleep pressure. Since *Lhx6*-positive ZI neurons are essential for the accumulation of induced sleep pressure (Liu et al. 2017a; Lee et al. 2025), and selective disruption of *Lhx6*-positive ZI neuron development by the loss of function of *Nkx2-2* severely disrupts sleep pressure-induced Fos expression in *Lhx6*-negative neurons, this implies that *Lhx6*-positive neurons coordinate these broader sleep pressure-dependent changes in activity across the ZI. This conclusion is further supported by the extensive reciprocal connections that exist among ZI neurons (Londei et al. 2024). In the case of induced sleep pressure, this may reflect the fact that glutamatergic neurons of the thalamic reuniens nucleus selectively project to *Lhx6*-positive ZI neurons and show sleep pressure-dependent increases in synaptic strength. However, this cannot account for the effects of naturally induced changes in sleep pressure, as this does not induce changes in the activity of reuniens neurons (Lee et al. 2025).

### Open questions and future directions

This raises the broader question of exactly how *Lhx6*-positive ZI neurons are selectively responsive to naturally occurring changes in sleep pressure. Our findings do not identify a molecularly distinct neuronal subpopulation that is responsive to any type of sleep pressure change, although *Calb2*-positive cells/*Lhx6*-positive cells appear to be less responsive. SnRNA-Seq from *Fos*-trapped neurons in the thalamic reuniens nucleus failed to identify any other molecular markers of specific neuronal subtypes responsive to induced sleep pressure (Lee et al. 2025), and the response properties of individual neurons in the ZI may likewise be determined by connectivity patterns that do not generally correlate with their gene expression profiles. *Lhx6*-positive ZI neurons may selectively receive synaptic input from an as yet unidentified neuronal subpopulation that is activated by naturally occurring changes in sleep pressure, or may sense these changes directly through as yet uncharacterized mechanisms. A more detailed analysis of the presynaptic inputs to these neurons that builds on previous work using rabies-based viral tracers (Liu et al. 2017a) and analysis of sleep pressure-induced transcriptomic changes will help address this.

The divergent sleep phenotypes seen in mutants that selectively disrupt the development of *Lhx6*-positive ZI neurons imply a more complex role for the ZI in regulating sleep pressure than has been previously hypothesized. Loss of function of *Lhx6* in early hypothalamic neuroepithelium leads to a complete loss of *Lhx6*-positive ZI neurons that likely results from selective apoptosis (Kim et al. 2021a), and in turn leads to increased wake and decreased sleep (Liu et al. 2017a). In contrast, loss of function of *Nkx2-2* in *Lhx6*-positive neural precursors reduces but does not completely eliminate *Lhx6*-positive ZI neurons, instead broadly disrupting the development and distribution of multiple subtypes of ZI neurons. This both dramatically reduces the induction of *Fos* that is induced by both naturally occurring and induced sleep pressure, and leads to significantly increased sleep. This implies that while *Lhx6*-positive ZI neurons are essential for sensing and signaling appropriate levels of sleep pressure both to *Lhx6*-negative ZI neurons and to arousal-promoting neurons in other brain regions, broad disruptions in the composition and organization of neuronal subtypes within the ZI like those that result from *Nkx2-2* loss of function may instead result in excessive activation of inhibitory projections to arousal-promoting neurons. This, in turn, underscores the importance of connectivity among specific neuronal populations of the ZI in maintaining appropriate levels of sleep homeostasis. A systematic *in vivo* analysis of changes in the activity and connectivity of both *Lhx6*-positive and *Lhx6*-negative neurons of the ZI will help clarify the dynamic regulation of these neural circuits, sensing, and signaling changes in sleep pressure.

## Materials and Methods

### Mouse colony maintenance

All experimental animal procedures were approved by the Johns Hopkins University Institutional Animal Care and Use Committee. All mice were housed in a climate-controlled facility (14-h light and 10-h dark cycle) with ad libitum access to food and water.

C57BL/6 were ordered from the Jackson Laboratory. Lhx6-iCre (B6;CBA-Tg(Lhx6-icre)1Kess/J, JAX#026555) and *Nkx2-2^lox/lox^* (Mastracci, Lin, and Sussel 2013) were crossed to generate *Lhx6-Cre;Nkx2-2^lox/l+^* mice. *Lhx6-Cre;Nkx2-2^lox/l+^* mice were then bred to generate *Lhx6-Cre;Nkx2-2^lox/l+^*or *Lhx6-Cre;Nkx2-2^lox/lox^* mice.

### Sleep deprivation

Between 8-12 weeks of age, mice were moved to cages in a climate-controlled room, given food and water ad libitum at 4:30 AM, and allowed an hour and a half to acclimate to the new environment before experimentation. Sleep deprivation began at 6:30 AM (ZT0) and was facilitated by gentle brushing until 12:30 PM for a total of 6 hours. Control mice were undisturbed and allowed to sleep freely. Mice were sacrificed after six hours (ZT6) using cervical dislocation, corresponding to the collection of sleep deprivation and control groups, and after one, three, six and eight hours of recovery sleep; corresponding to seven hours (ZT7), nine hours (ZT9), twelve hours (ZT12) and fourteen hours (ZT14) after lights on, respectively. Brains were collected within 15 minutes of sacrifice and stored in OCT at -80°C until sectioning.

### Tissue sectioning

Sections were coronally sliced between 10 and 14 μm using Leica Cryotstat CM3050S and placed on Superfrost Plus Microscope Slides or Xenium Slides. For Hiplex analysis, sections were collected every 30-42 µm. For Xenium analysis, sections were collected every 100 µm.

### Hiplex analysis

Multiplex in situ hybridization was performed according to the fresh-frozen protocol listed by the RNAscope HiPlex v2 assay (ACDBio) using the probes (also provided by ACDBio) found in Table 1.

### Xenium analysis

Xenium *In Situ* Gene expression was performed according to the fresh-frozen protocol listed by 10x Genomics (Ma et al. 2024).

### Imaging

Slides were imaged using the Olympus widefield microscope at 20x, 40x, and 60x Oil magnifications. Images were captured using z-stacked montages using DAPI, GFP, TRITC, Cy5, and Cy7 filters. Images were further processed using Slidebook Reader (Intelligent Imaging Innovations, Denver, CO, USA) and ImageJ (Schneider, Rasband, and Eliceiri 2012; Blattner et al. 2014; Chalfoun et al. 2017).

### Hiplex: Cell counting, segmentation & data assembly

Cells were segmented, counted, and assigned puncta using pipelines developed in Cell Profiler (Stirling et al. 2021). From these assignments, cells were further categorized into high, low, and negative expressing cell types based on the number of puncta expressed per probe (Table 2). With this information, a master sheet was created describing each cell, its assigned puncta, experimental group, and cell type (Supplemental Dataset 1).

### Xenium: Analysis, Segmentation & Visualization

Data visualization was performed using the Xenium Explorer 1.3 (Ma et al. 2024) and the Seurat package in R (Satija et al. 2015; Hao et al. 2024).

### Piezo analysis of sleep/wake behavior

The PiezoSleep system (Signal Solutions, LLC) was used to record and analyze sleep of mice under the light/dark (12h/12h) condition as described previously (Yaghouby et al. 2016). Briefly, Piezoelectric motion sensor film detects the sleep and wakefulness states of individual mice. The signal was amplified via a PiezoSleep in-line amplifier and collected via a Calamari SAS (8-channel) data acquisition system connected to a computer with PiezoSleep v.2.18 software installed. Sleep amount and bout duration were analyzed offline using SleepStats v.4.

### Statistical analysis

Paired t-tests, Fisher’s exact tests, one-way ANOVAs, two-way ANOVAs, Kruskal-Wallis tests (in cases of non-parametric data), mixed-effect analysis, Tukey’s Multiple Comparison tests, and Dunn’s Multiple Comparison tests were performed using GraphPad Prism version 10.0.0 for Windows (GraphPad Software, Boston, Massachusetts, USA). The Seurat “FindAllMarkers” function with assay “SCT” and default parameters was used for analyzing differential gene expression, using the number of total mRNAs and genes as a variable. All bar graphs show mean and standard deviation (SD), with individual data points plotted. Significance was determined using an alpha of 0.05.

## Acknowledgements

We thank W. Yap for comments on the manuscript. This work was supported by a National Science Foundation graduate fellowship to P.W.C and a grant from the NIH (R01MH126676) to S.B.

## Resource Availability

## Lead contact

Further information and requests for resources and reagents should be directed to and will be fulfilled by the Lead Contact, Seth Blackshaw.

## Materials availability

All unique/stable reagents generated in this study are available from the Lead Contact upon request.

## Supplemental Dataset 1: Hiplex data for all samples

The dataset contains all of the cells identified in the zona incerta across all experimental conditions and genotypes. Each cell is delineated by a unique cell ID name and contains corresponding information regarding experimental group, genotype, location, markers expressed, and cell type.

---

**Supplementary Figure 1:**
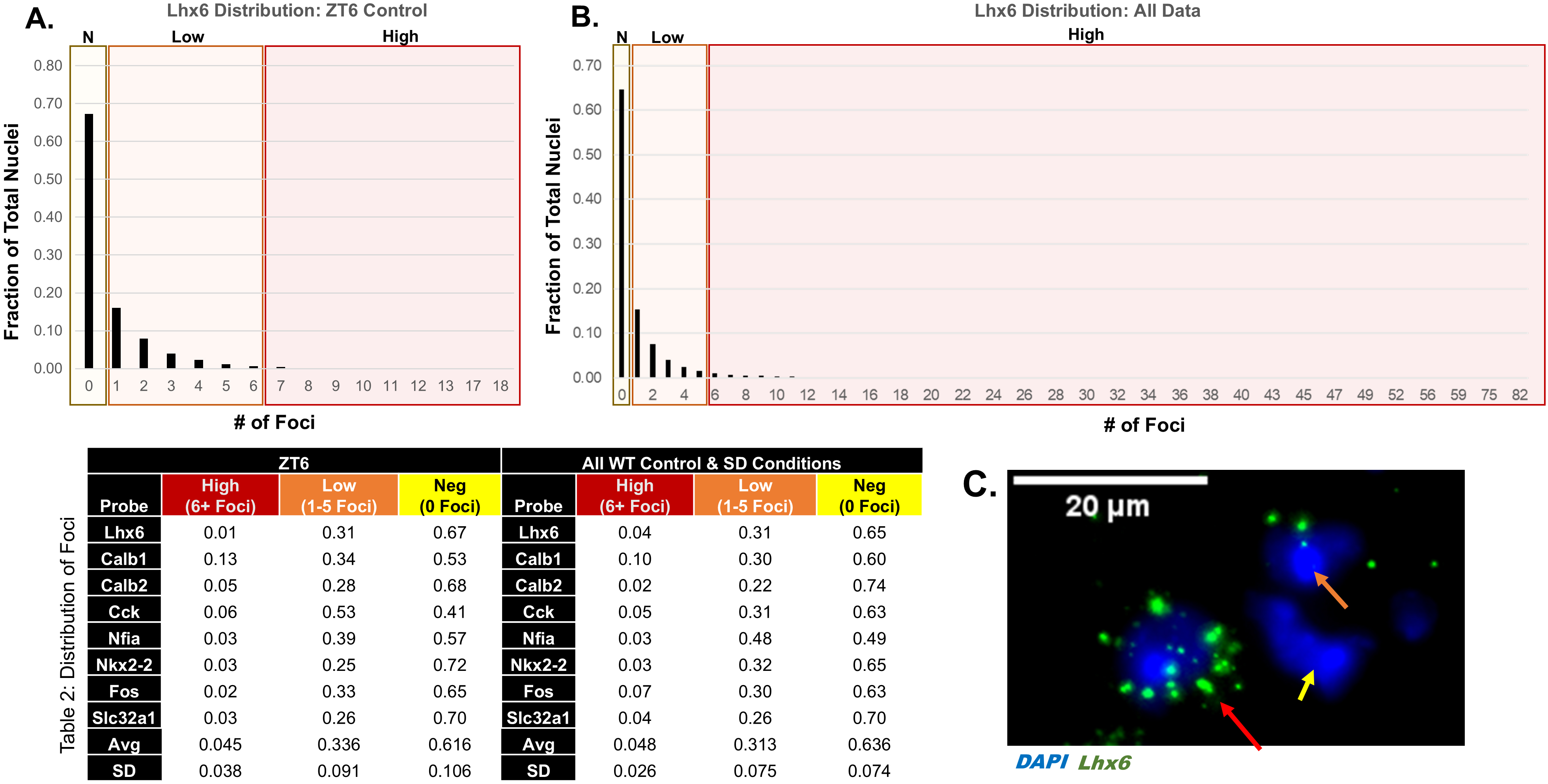
Differential cellular expression levels of *Lhx6* and other cell subtype-specific molecular markers in zona incerta. **A**. Bar graph depicts the fraction of cells that express up to 18 *Lhx6* puncta (green) in ZT6 controls. The graph is further binned by negative cells (0 puncta expressed, yellow), low-expressing cells (1-5 puncta, orange), and high-expressing cells (≥ 6 puncta, red). **B**. Bar graph depicts the fraction of cells that express up to 82 *Lhx6* puncta across all conditions in wild-type mice. The graph is further binned by negative cells (0 puncta expressed, yellow), low-expressing cells (1-5 puncta, orange), and high-expressing cells (≥ 6 puncta, red). **C**. In situ hybridization exhibiting examples of high expressing Lhx6 cells (red arrow), low expressing Lhx6 cells (orange arrow), and Lhx6 negative cells (yellow arrow). Scale bar=20 µm.

**Supplementary Figure 2:**
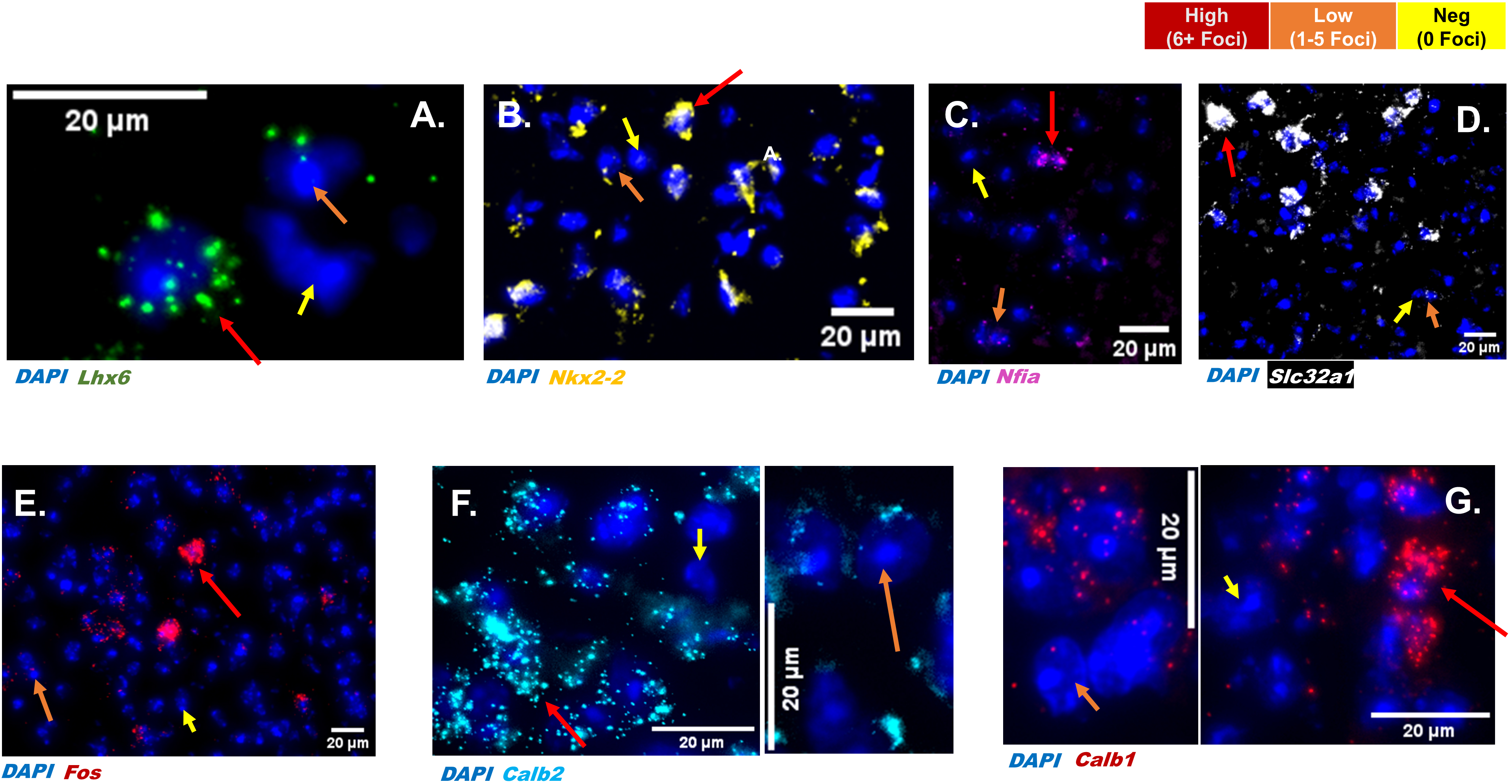
Examples of cells expressing high and low levels of cell subtype-specific molecular markers. *In situ* hybridization exhibiting examples of negative cells (yellow arrow), low-expressing cells (orange arrow), and high-expressing cells (red arrow) for *Lhx6* (green, **A**), *Nkx2-2* (yellow, **B**), *Nfia* (purple, **C**), *Slc32a1* (white, **D**), *Fos* (red, **E**), *Calb2* (light blue, **F**), and *Calb1* (red, **G**).

**Supplementary Figure 3:**
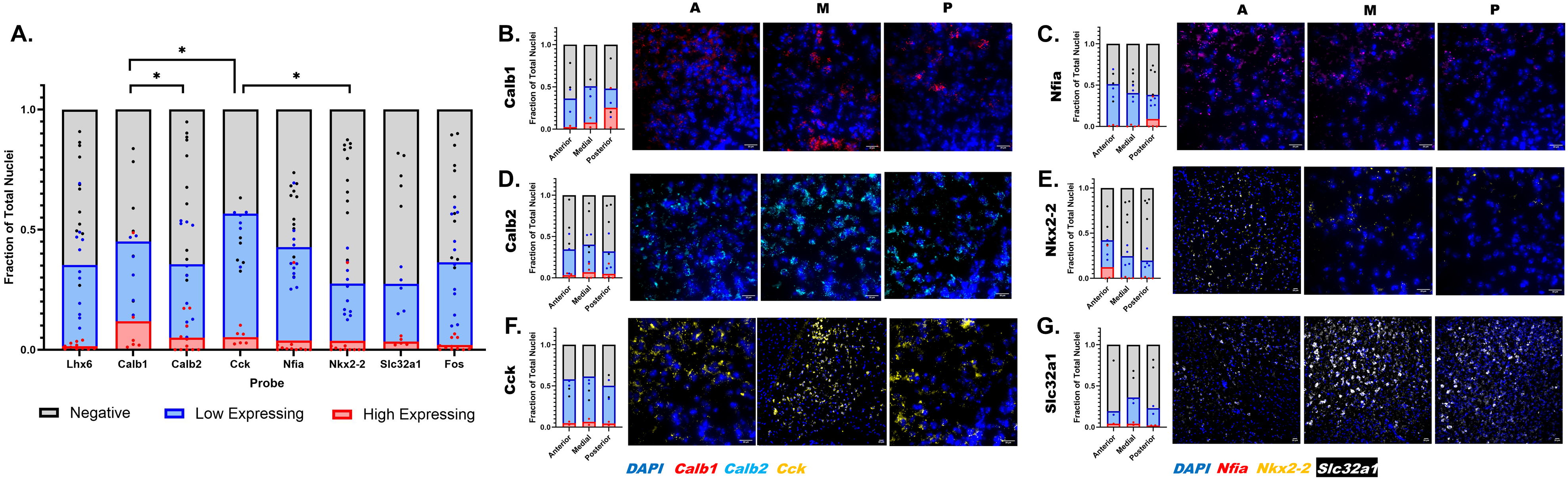
*Cck*- and *Calb1*-positive cells are the most abundant cell subtype-specific molecular markers examined in the zona incerta. **A**. Bar graph depicting the fraction of marker-positive cells at ZT6. *Lhx6*-positive cells comprise 35.2% of total cells in the ZI (high expressing [red] compromising 1.5±1.3% of cells, low expressing cells [blue] Lhx6 comprising 33.7±18.7%); *Calb1* comprise 45.1% (high:11.9±18.6%, low: 33.2±13.7%), *Calb2* comprise 35.6% (high: 5.0%±6.8%, low: 30.5±19.7%), *Cck* comprise 56.6% (high: 5.3±3.1%, low:51.3±8.7% ), *Nfia* 42.7% (high:3.9±10.8%, low: 38.8±12.8%), *Nkx2-2* 27.5% (high:3.7±10.8%, low: 23.8±10.3% ), *Slc32a1* 27.5% (high: 3.4±1.6%,low: 24.1±8.4%), *Fos* 36.4% (high: 2.0±2.4%,low: 34.4±18.4%). Significant differences in the abundance of low-expressing cells are observed between *Calb1* and *Calb2* (*p* = 0.0462, alpha=0.05), *Calb1* and *Cck* (*p* = 0.0336, alpha=0.05), and *Cck* and *Nkx2-2* (*p* = 0.0252, alpha=0.05). *Cck*, N = 6, Slc32a1, N = 5, all other probes N = 11.

**Supplementary Figure 4:**
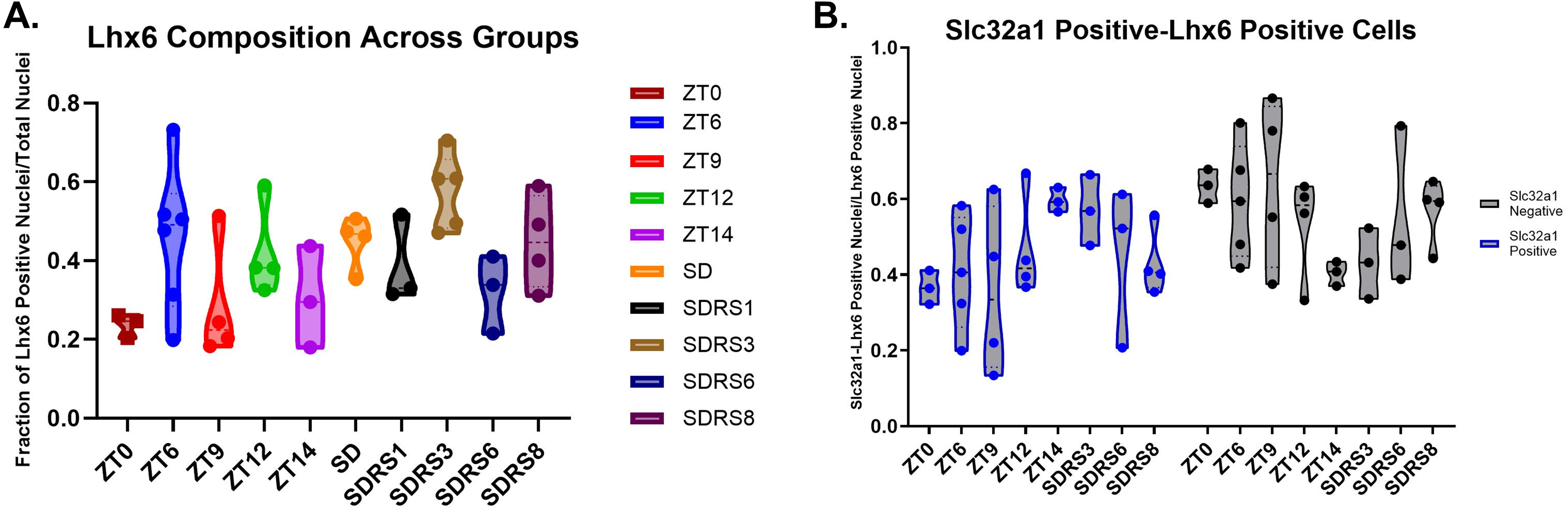
Number of GABAergic *Lhx6*-positive neurons remains constant across experimental groups. A Bar graph depicting the fraction of *Lhx6*-positive cells within each experimental group. No significant differences were found. B Violin plot depicting the fraction of *Slc32a1/Lhx6-*positive and *Slc32a1*-negative/*Lhx6*-positive cells amongst all *Lhx6*-positive cells within each experimental group.

**Supplementary Figure 5:**
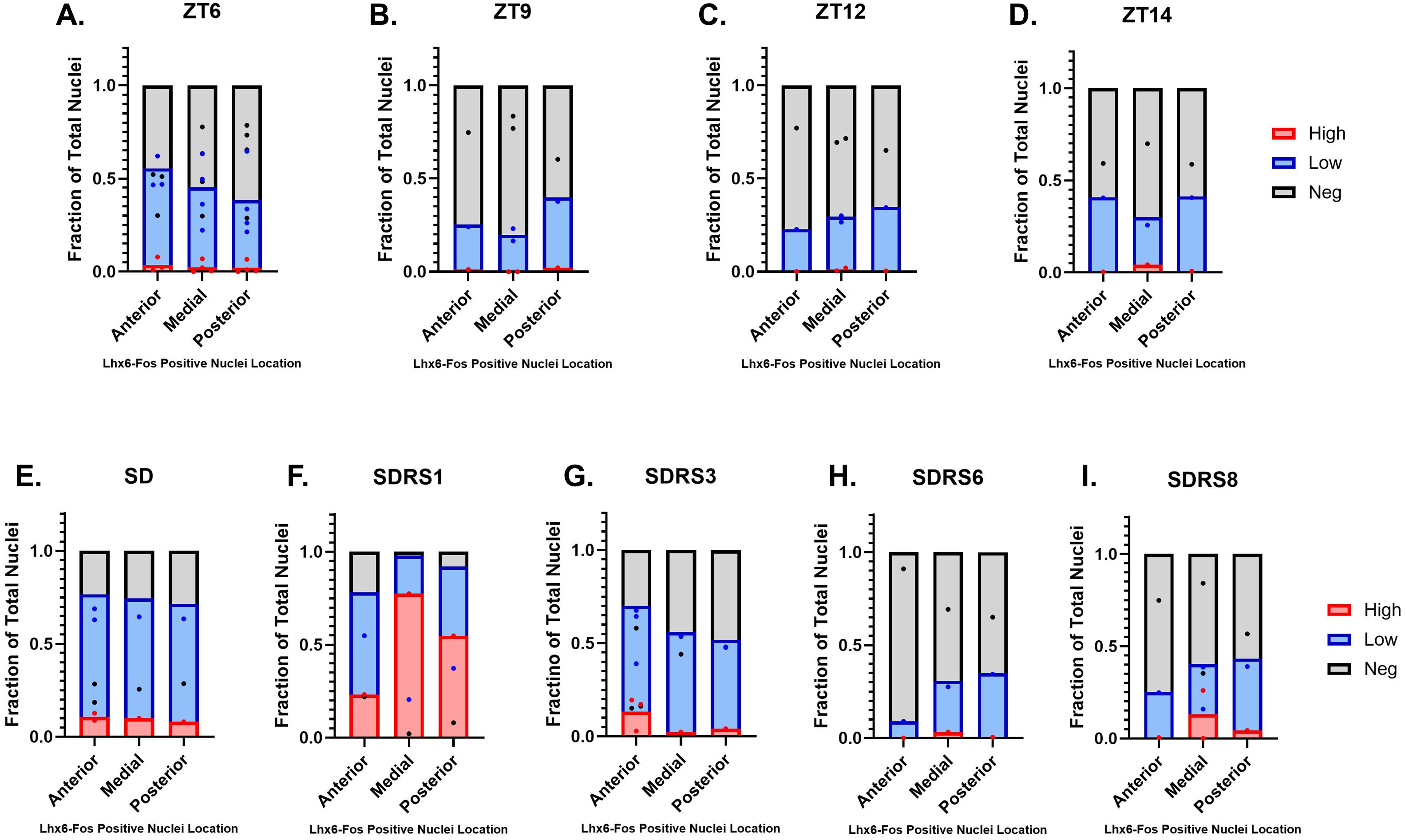
Differential sleep pressure-dependent *Fos* induction along the anterior-posterior axis of the zona incerta. **A-I.** Bar graphs depict *Fos* expression in low-expressing (blue) and high-expressing (red) *Lhx6*-positive cells as a fraction of total *Lhx6*-positive cells across the anterior to posterior axis for each experimental group. **A** ZT6 Anterior: 55.5% (high: 3.7±3.7%, low: 51.8±8.8%, N = 3), Medial: 45.2% (high: 2.4±3.2% , low:42.9±17.6% ,N = 4), Posterior: 38.5% (high: 2±3.1%, low: 36.5±19.4%, N = 4). **B** ZT9 Anterior: 25.3% (high: 1.2%, low: 24.1%, N = 1), Medial: 19.8% (high: 0%, low: 19.8±4.7%, N = 2), Posterior: 39.8% (high:2.1%, low: 37.7%, N = 1). **C** ZT12 Anterior: 22.8% (high: 0.1%, low: 22.7%, N = 1), Medial: 29.6% (high: 1.3±1.1%, low:28.3±2.4%, n=2), Posterior:34.9% (high: 0.4%, low: 34.5%, N = 1). **D** ZT14 Anterior: 40.8%, (high: 0.3%, low:40.5%, N = 1), Medial: 30% (high: 4.3%, low: 25.7%, N = 1), Posterior: 41.3% (high: 0.7%, low:40.6%, N = 1) **E** SD Anterior: 76.6% (high: 10.8±2.8%, low: 65.9±4.2%, N = 2), Medial: 74.4% (high: 9.9%, low: 64.5%, N = 1), Posterior 71.4% (high: 8%, low: 63.4%, N = 1). **F** SDRS1 Anterior: 78.1% (high: 23.2%, low: 54.9%, N = 1), Medial: 97.9% (high:77.4%, low: 20.5%, N = 1), Posterior: 92.1% (high:54.8%, low: 37.3%, N = 1) **G** SDRS3 Anterior: 70.2% (high: 13.2%±9%, low: 57%±15.7%, N = 3), Medial: 55.9% (high: 2.3%, low: 53.6%, N = 1),l Posterior: 51.9% (high: 4.1%, low: 47.8%, N = 1) **H** SDRS6 Anterior: 9% (high: 0%, low: 9%, N = 1), Medial: 30.8% (high: 3.2%, low: 27.6%, N = 1), Posterior: 34.9% (high: 0.5%, low: 34.4%, N = 1) **I** SDRS8 Anterior: 25.2% (high:0.4%, low: 24.8%, n=1), Medial: 40.3% (high:26±26%, low:27.3±16.1%, N = 2), Posterior: 43.3% (high:4.3%, low: 39%, N = 1).

**Supplementary Figure 6:**
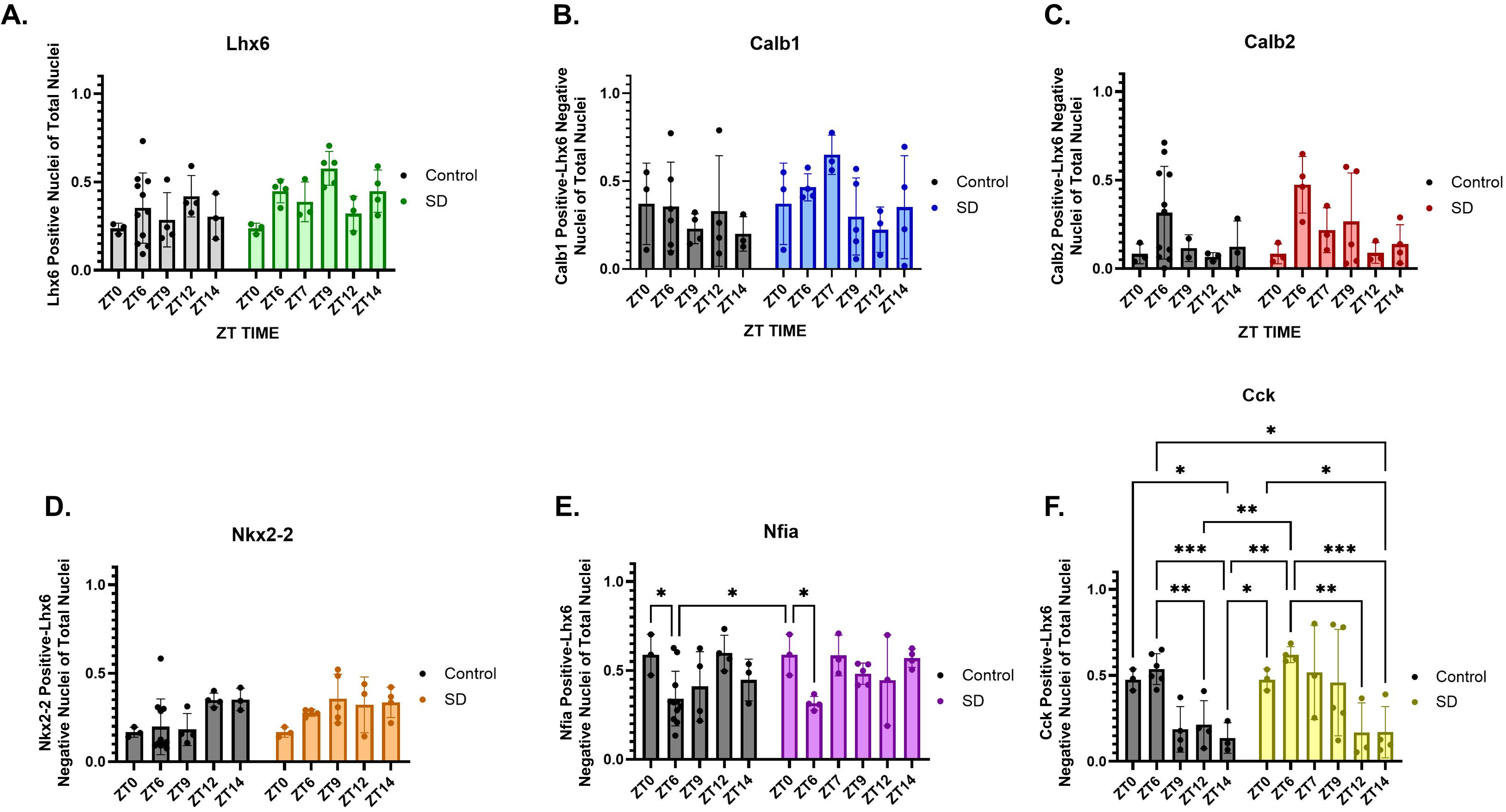
*Cck*, but not other cell subtype-specific markers, shows altered expression in response to changes in sleep pressure. Bar graphs depicting the relative fraction of zona incerta cells expressing the indicated markers, including **A**. *Lhx6* (green), **B**. *Calb1* (blue), **C.** *Calb2* (red), **D**. *Nkx2-2* (orange), and **E**. *Nfia* (purple). Significant differences: Control ZT0 v. Control ZT6 (*p* = 0.0190), SD ZT0 v SD ZT6 (*p* = 0.0190). **F.** *Cck* (yellow). Significant differences: ZT6:Control vs. ZT12:Control (*p* = 0.0026), ZT6:Control vs. ZT14:Control (*p* = 0.0005), ZT0:SD vs. ZT14:SD (*p* = 0.0474), ZT6:SD vs. ZT12:SD (*p* = 0.0026), ZT6:SD vs. ZT14:SD (*p* = 0.0005).

**Supplementary Figure 7:**
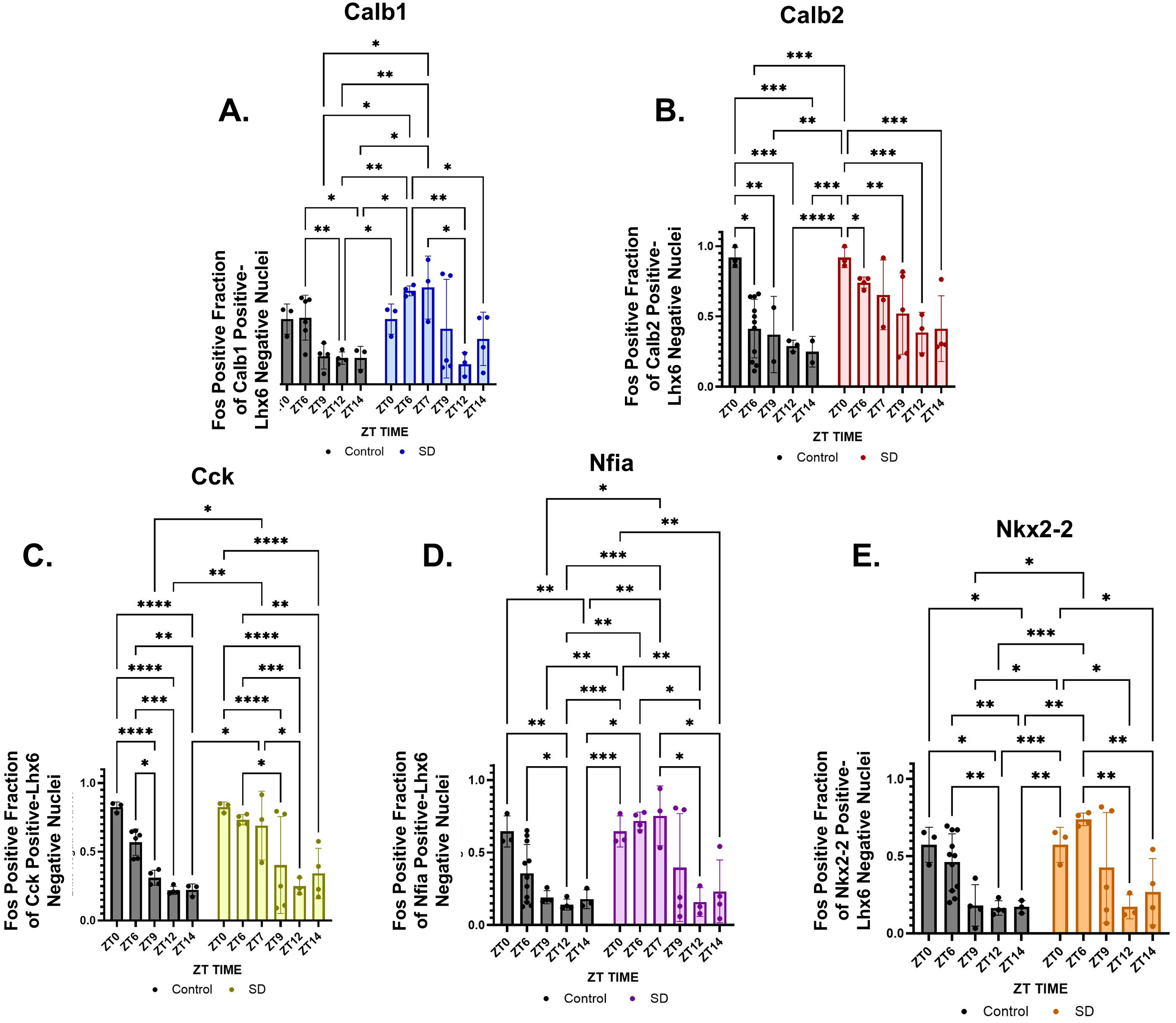
*Lhx6*-negative/*Calb2*-positive and *Lhx6*-negative/*Cck*-positive cells are less responsive to experimentally induced sleep pressure. Bar graphs depicting the *Fos*-positive fraction of marker-positive, *Lhx6*-negative cells under naturally occurring (grey) and induced sleep pressure (colored).Significant differences: **A.** *Calb1* (blue): ZT6:Control vs. ZT12:Control (*p* = 0.0030), ZT6:Control vs. ZT14:Control (*p* = 0.0420), ZT6:SD vs. ZT12:SD(*p* = 0.0030), ZT6:SD vs. ZT14:SD(*p* = 0.0420), ZT7:SD vs. ZT9:Control(*p* = 0.0291) , ZT7:SD vs. ZT12:Control (*p* = 0.0020), ZT7:SD vs. ZT12:SD (*p* = 0.0279), ZT7:SD vs. ZT14:Control (*p* = 0.0155). **B.** *Calb2* (red): ZT0:Control vs. ZT6:Control (*p* = 0.0139), ZT0:Control vs. ZT9:Control (*p* = 0.0042), ZT0:Control vs. ZT12:Control (*p* = 0.0004), ZT0:Control vs. ZT14:Control (*p* = 0.0004), ZT0:SD vs. ZT6:SD (*p* = 0.0139), ZT0:SD vs. ZT9:SD(*p* = 0.0042), ZT0:SD vs. ZT12:SD (*p* = 0.0004), ZT0:SD vs. ZT14:SD (*p* =0.0004). **C**. *Cck* (yellow) ZT0:Control vs. ZT9:Control (*p* < 0.0001), ZT0:Control vs. ZT12:Control (*p* < 0.0001), ZT0:Control vs. ZT14:Control (*p* < 0.0001), ZT0:SD vs. ZT9:SD (p < 0.0001), ZT0:SD vs. ZT12:SD (p<0.0001), ZT0:SD vs. ZT14:SD (*p* < 0.0001), ZT6:Control vs. ZT9:Control (*p* = 0.0136), ZT6:Control vs. ZT12:Control (*p* = 0.0003), ZT6:Control vs. ZT14:Control (*p* = 0.0020), ZT6:SD vs. ZT9:SD (*p* = 0.0136), ZT6:SD vs. ZT12:SD (*p* = 0.0003), ZT6:SD vs. ZT14:SD (*p* = 0.0020), ZT7:SD vs. ZT9:Control (*p* = 0.0447), ZT7:SD vs. ZT12:Control (*p* = 0.0029), ZT7:SD vs. ZT12:SD (*p* = 0.0307), ZT7:SD vs. ZT14:Control (*p* = 0.01100). **D.** *Nfia* (purple): ZT0:Control vs. ZT12:Control (*p* = 0.0028), ZT0:Control vs. ZT14:Control (p=0.0076), ZT0:SD vs. ZT12:SD (*p* = 0.0028), ZT0:SD vs. ZT14:SD (*p* = 0.0076), ZT6:Control vs. ZT12:Control (*p* = 0.0259), ZT6:SD vs. ZT12:SD (*p* = 0.0259), ZT7:SD vs. ZT9:Control (*p* = 0.0107), ZT7:SD vs. ZT12:Control (*p* = 0.0007), ZT7:SD vs. ZT12:SD (*p* = 0.0260), ZT7:SD vs. ZT14:Control (*p* = 0.0020), ZT7:SD vs. ZT14:SD (*p* = 0.0460)., **E.** *Nkx2-2* (orange): ZT0:Control vs. ZT12:Control (*p* = 0.0127), ZT0:Control vs. ZT14:Control (*p* = 0.0336), ZT0:SD vs. ZT12:SD (*p* = 0.0127), ZT0:SD vs. ZT14:SD (*p* = 0.0336), ZT6:Control vs. ZT12:Control (*p* = 0.0014), ZT6:Control vs. ZT14:Control (*p* = 0.0062), ZT6:SD vs. ZT9:Control(*p* = 0.0128), ZT6:SD vs. ZT9:SD (*p* = 0.0004), ZT6:SD vs. ZT12:Control (*p* = 0.0004), ZT6:SD vs. ZT12:SD (*p* = 0.0014), ZT6:SD vs. ZT14:Control (*p* = 0.0022), ZT6:SD vs. ZT14:SD (*p* = 0.0062).

**Supplementary Figure 8:**
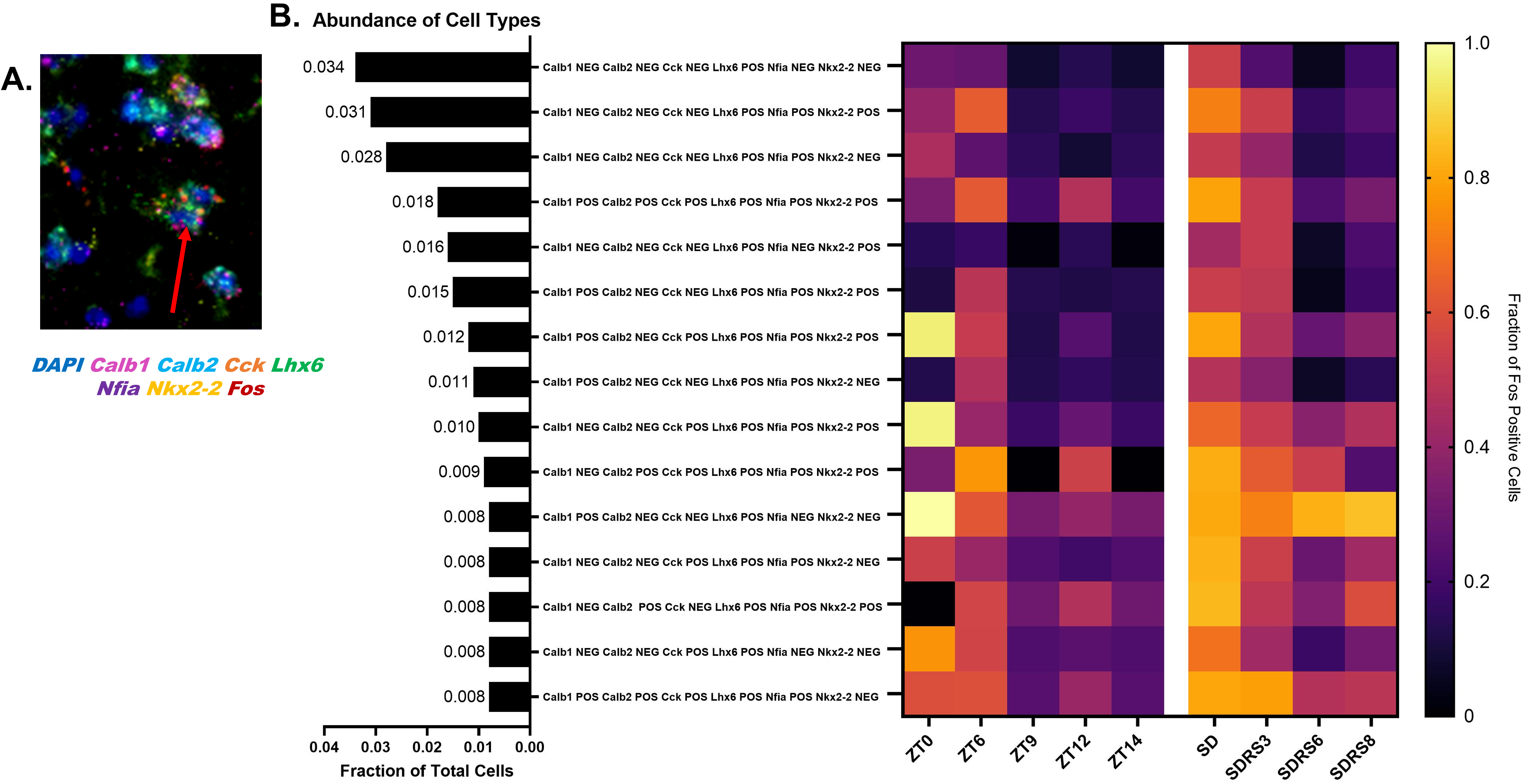
*Fos* induction is sustained for up to 8 hours following experimentally induced sleep pressure in *Lhx6*-positive cells. **A.** *In situ* hybridization identifying example of *Calb1/Calb2/Cck/Lhx6/Nfia/Nkx2-2/Fos* positive cells (cell type identified in row four of panel B). **B.** Bar graph (left) and heatmap (right) depicting the abundance of Hiplex-identified, *Lhx6*-positive cell types in wildtype controls and their expression under natural circadian conditions and induced sleep deprivation.

**Supplementary Figure 9:**
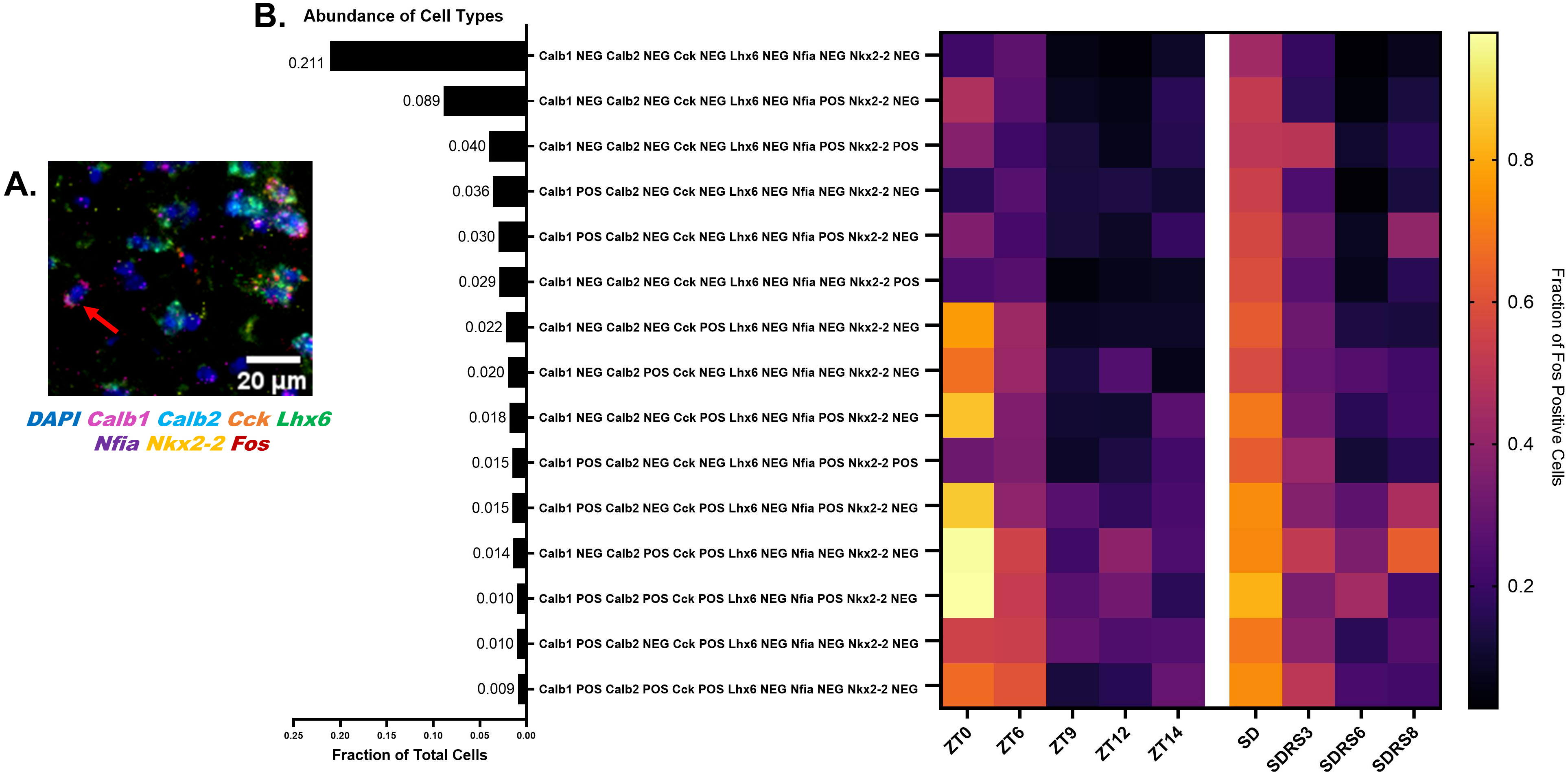
*Calb1*-positive and *Cck*-positive/*Lhx6*-negative cells maintain *Fos* expression in conditions of low sleep pressure. **A.** *In situ* hybridization for *Calb1*-positive cells. **B.** Bar graph (left) and heatmap (right) depicting the abundance of HiPlex-identified, *Lhx6*-negative cell subtypes in wildtype controls and their expression under natural circadian conditions and experimentally-induced sleep deprivation.

**Supplementary Figure 10:**
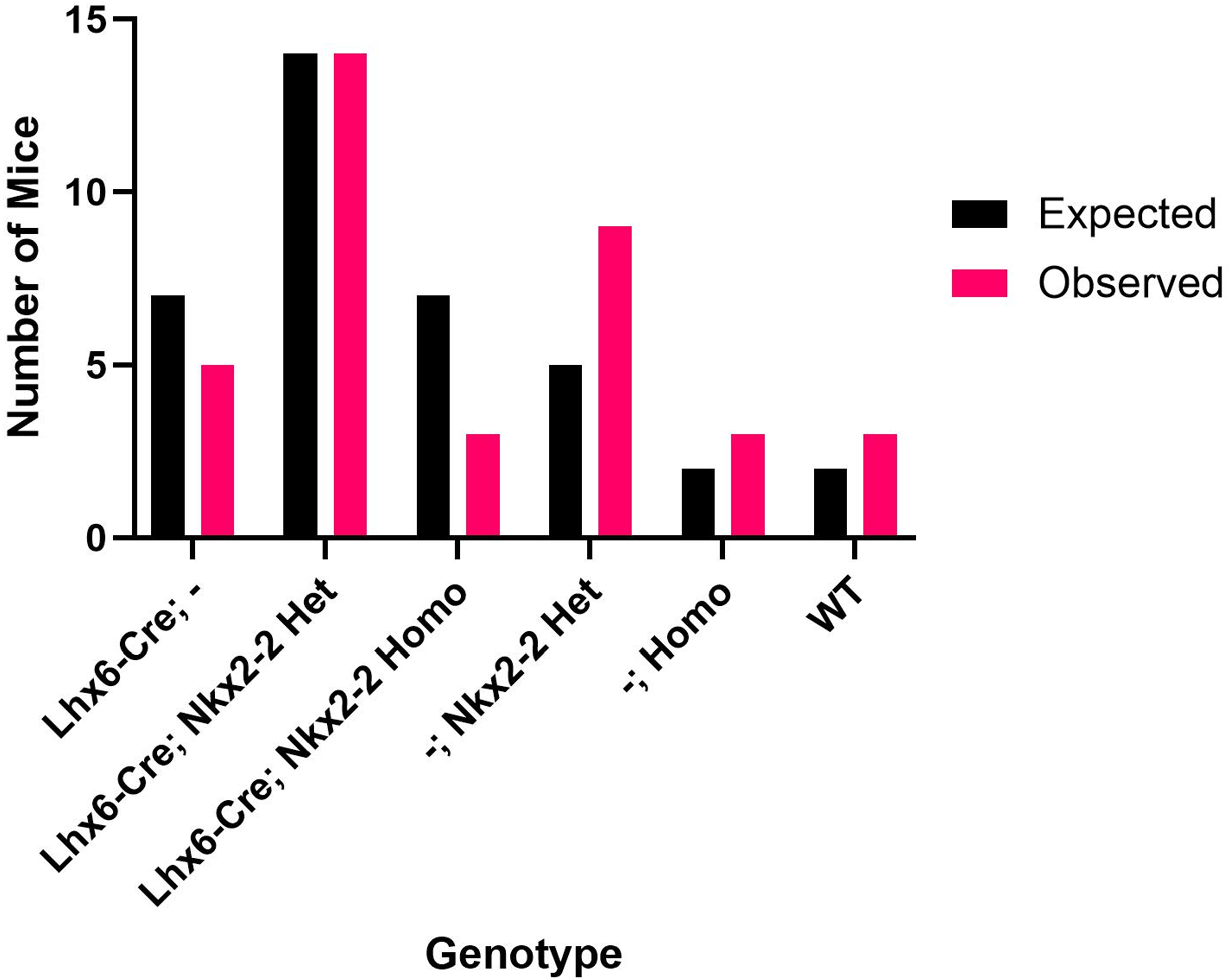
*Lhx6-Cre;Nkx2-2^lox/lox^* mice are born at expected Mendelian ratios. Bar graph depicting the expected and observed number of live births per genotype from a *Lhx6-Cre;Nkx2-2^lox/lox^* x *Lhx6-Cre;Nkx2-2^lox/lox^* cross. The animals used for this graph totaled 37 animals across 4 liters.

**Supplementary Figure 11:**
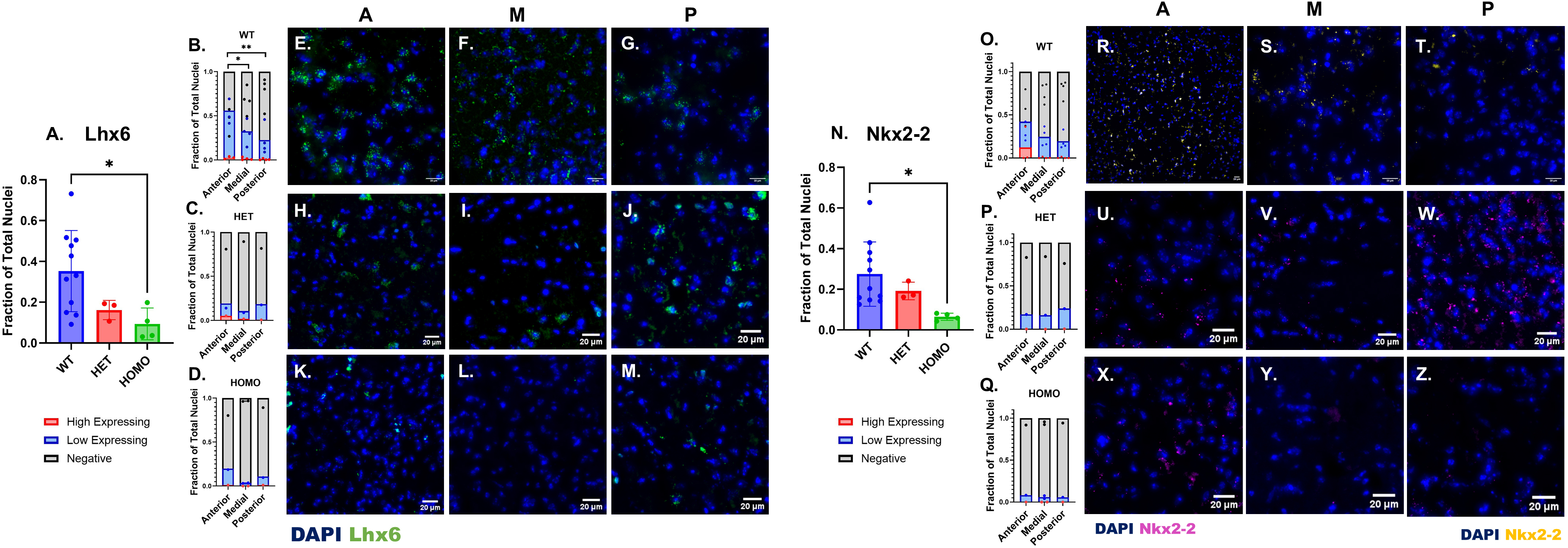
*Nkx2-2* mutants show reduced expression of *Lhx6* and *Nkx2-2* in the zona incerta and redistribution of both markers along the anterior-posterior axis. **A** Bar graph depicting the fraction of Lhx6-positive cells of total cells amongst experimental groups: wildtype (WT, blue), *Lhx6-Cre;Nkx2-2^lox/+^* (HET, red), *Lhx6-Cre;Nkx2-2^lox/lox^* (HOMO, green). Significant difference found between wildtype and *Lhx6-Cre;Nkx2-2^lox/lox^* (HOMO) mice (p = 0.0449). **B** Bar graph depicting fraction of Lhx6 positive cells (high expressing: red, low expressing: blue) of total cells along the anterior to posterior axis in wildtype (WT) mice at ZT6. Significant differences were found between the low-expressing cells at the anterior and medial (*p* = 0.0187, alpha = 0.05), and the anterior and posterior positions (*p* = 0.0012, alpha=0.05). **C** Bar graph depicting the fraction of *Lhx6*-positive cells (high expressing: red, low expressing: blue) of total cells along the anterior to posterior axis in *Lhx6-Cre;Nkx2-2^lox/+^* (HET) mice at ZT6. **D.** Bar graph depicting the fraction of *Lhx6*-positive cells (high expressing: red, low expressing: blue) of total cells along the anterior to posterior axis in *Lhx6-Cre;Nkx2-2^lox/lox^* (HOMO) mice at ZT6. **E-G.** *In situ* hybridization depicting *Lhx6* expression (green) along the anterior-posterior axis (AMP) in wild-type mice. Scale bar = 20 µm. **H-J.** *In situ* hybridization depicting *Lhx6* expression (green) along the anterior-posterior axis (AMP) in *Lhx6-Cre;Nkx2-2^lox/+^* (HET) type mice. 20 µm scale bars. **K-M.** *In situ* hybridization depicting *Lhx6* expression (green) along the anterior-posterior axis (AMP) in *Lhx6-Cre;Nkx2-2^lox/lox^*(HOMO) type mice. 20 µm scale bars. **N.** Bar graph depicting the fraction of *Nkx2-2*-positive cells of total cells amongst experimental groups: wildtype (WT, blue), *Lhx6-Cre;Nkx2-2^lox/+^*(HET, red), *Lhx6-Cre;Nkx2-2^lox/lox^*(HOMO, green). Significant difference found between wildtype and *Lhx6-Cre;Nkx2-2^lox/lox^* (HOMO) mice (*p* = 0.0368). **O.** Bar graph depicting fraction of *Nkx2-2*-positive cells (high expressing: red, low expressing: blue) of total cells along the anterior to posterior axis in wildtype (WT) mice at ZT6. **P.** Bar graph depicting the fraction of *Nkx2-2*-positive cells (high expressing: red, low expressing: blue) of total cells along the anterior to posterior axis in *Lhx6-Cre;Nkx2-2^lox/+^* (HET) mice at ZT6. **Q.** Bar graph depicting the fraction of *Nkx2-2*-positive cells (high expressing: red, low expressing: blue) of total cells along the anterior to posterior axis in *Lhx6-Cre;Nkx2-2^lox/lox^* (HOMO) mice at ZT6. **R-T.** *In situ* hybridization depicting *Nkx2-2* expression (yellow) along the anterior-posterior axis (AMP) in wildtype mice. 20 µm scale bars. **U-W.** *In situ* hybridization depicting *Nkx2-2* expression (purple) along the anterior-posterior axis (AMP) in *Lhx6-Cre;Nkx2- 2^lox/+^* (HET) type mice. 20 µm scale bars. **X-Z.** *In situ* hybridization depicting *Nkx2-2* expression (purple) along the anterior-posterior axis (AMP) in *Lhx6-Cre;Nkx2-2^lox/lox^* (HOMO) type mice. 20 µm scale bars.

**Supplementary Figure 12:**
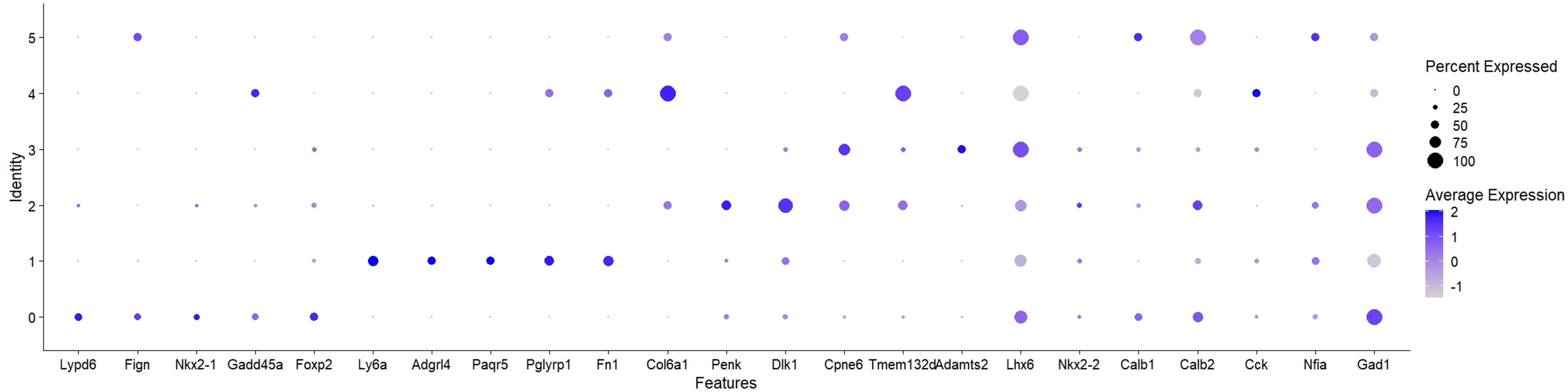
Xenium-based identification of additional markers of Lhx6-positive zona incerta neurons. A dot plot depicts the relative expression of individual Hiplex markers and markers identified by differential expression analysis.

**Supplementary Figure 13:**
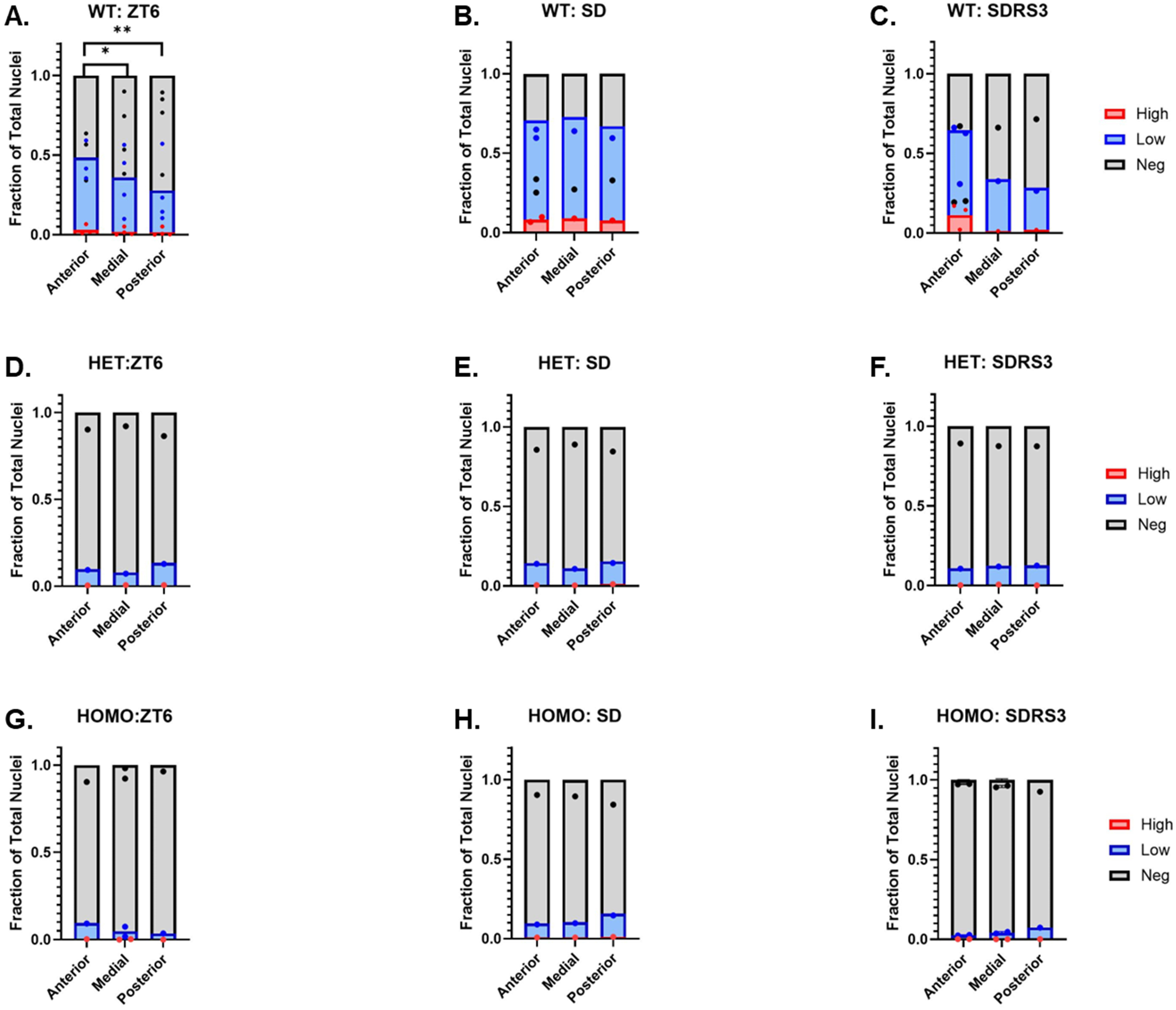
*Nkx2-2*-deficient mice show altered distribution of sleep deprivation-induced Fos expression along the anterior-posterior axis of the zona incerta. **A-C**. Fraction of *Fos*-positive (high expressing, red; low expressing, blue) cells in wildtype mice along the anterior-posterior axis at ZT6 (A), SD (B), and SDRS3 (C). D-F. Fraction of *Fos*-positive (high expressing, red; low expressing, blue) cells in *Lhx6-Cre;Nkx2-2^lox/+^* (HET) mice along the anterior-posterior axis at ZT6 (D), SD (E), and SDRS3 (F). G-I. Fraction of *Fos*-positive (high expressing, red; low expressing, blue) cells in *Lhx6-Cre;Nkx2-2^lox/lox^* (HOMO) mice along the anterior-posterior axis at ZT6 (G), SD (H), and SDRS3 (I).

**Supplementary Figure 14:**
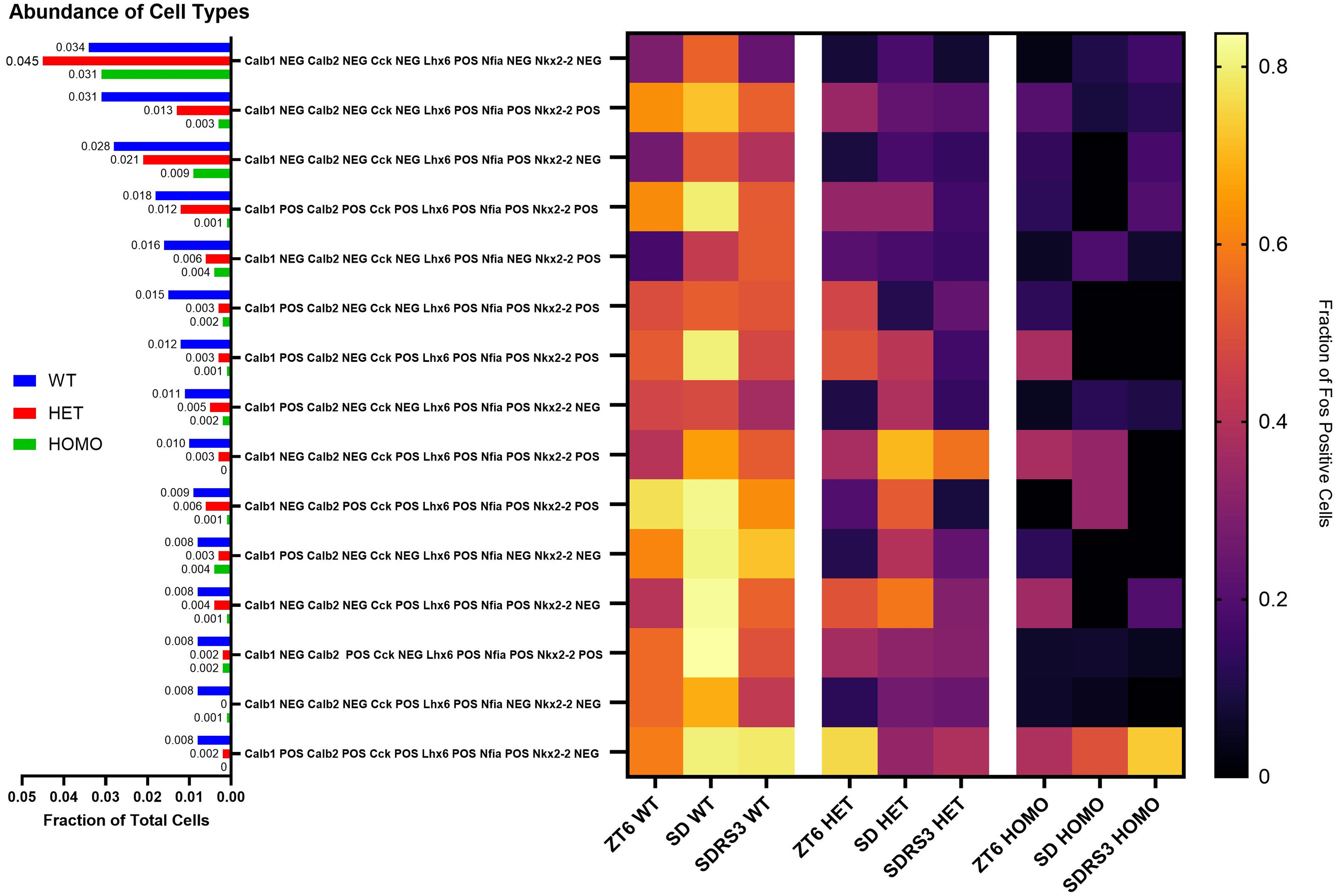
*Nkx2-2*-deficient mice show reduced numbers and defective Fos induction in Lhx6-positive neurons expressing subtype-specific markers. *Nkx2-2*-deficient mice show reduced numbers and defective Fos induction in Lhx6-positive neurons expressing subtype-specific markers. Bar graph (left) and heatmap (right) depicting the abundance of *Lhx6*-positive cells that are also positive for other cell subtype-specific markers in wildtype controls, *Nkx2-2* mutant knockouts, and the expression of these markers under natural circadian conditions and induced sleep deprivation.

**Supplementary Figure 15:**
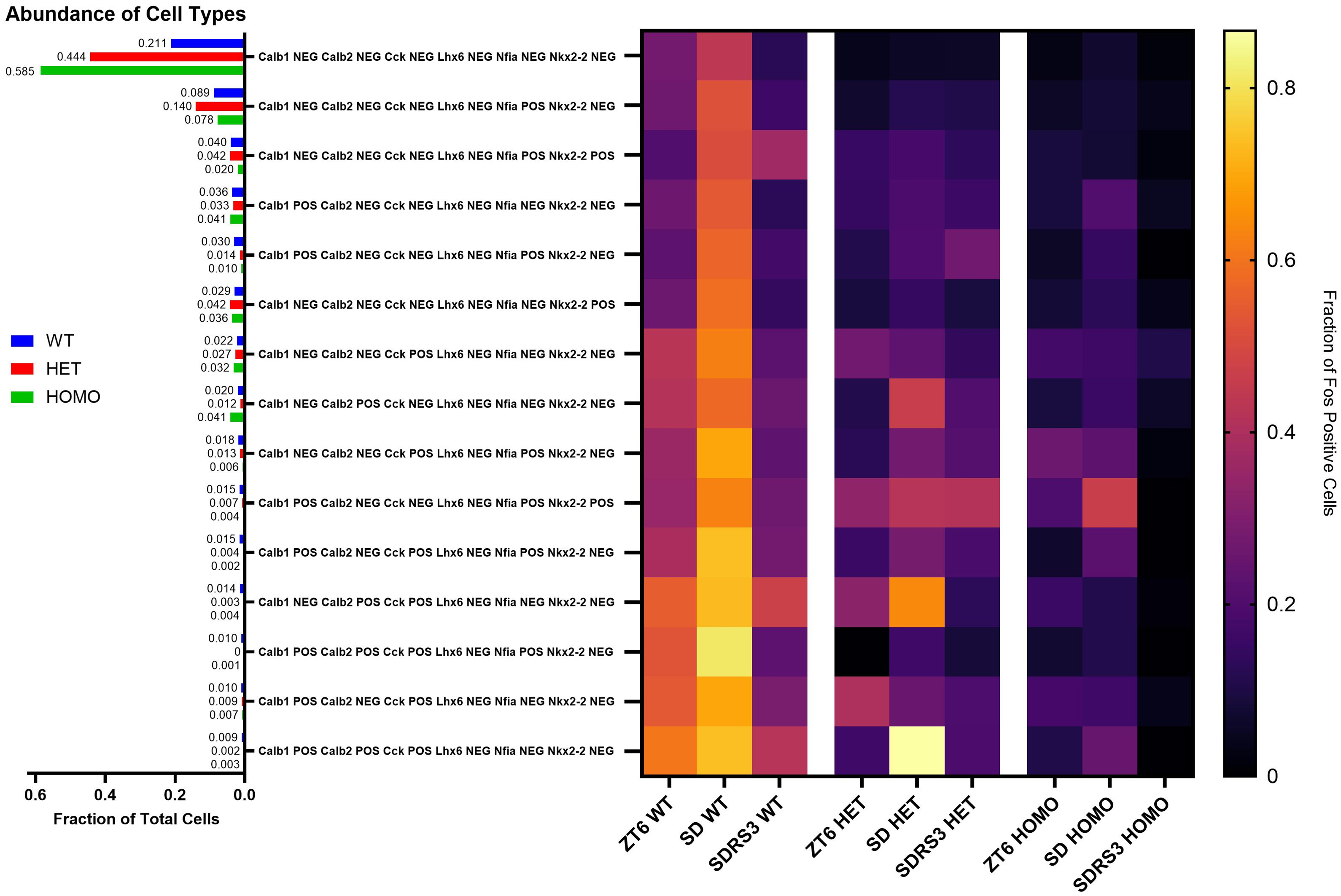
*Nkx2-2*-deficient mice maintain sleep deprivation-induced Fos expression in *Lhx6*-negative/*Calb1*-positive and *Lhx6*-negative/*Cck*-positive cells. *Nkx2-2*-deficient mice show reduced numbers and defective Fos induction in Lhx6-positive neurons expressing subtype-specific markers. Bar graph (left) and heatmap (right) depicting the abundance of Hiplex-identified *Lhx6*-negative cell subtypes in wildtype controls, *Nkx2-2* mutant knockouts, and their expression under natural circadian conditions and induced sleep deprivation.

